# Echoes as Signal, Not Noise: Reverberation Sharpens Sensitivity to Temporal Structure

**DOI:** 10.64898/2025.12.15.693365

**Authors:** Peirun Song, Lingling Zhang, Yuanhao Huang, Haoxuan Xu, Yuying Zhai, Xuehui Bao, Hangting Ye, Ishrat Mehmood, Nayaab Shahir Pandit, Yanyan Wang, Zhiyi Tu, Pei Chen, Tingting Zhang, Xuan Zhao, Xiongjie Yu

## Abstract

Sensitivity to temporal structure is fundamental to hearing, yet how natural reverberation shapes this sensitivity is unclear. Here we identify echo-facilitated temporal sensitivity (EFTS), a principle whereby the auditory system exploits echoes to selectively enhance responses to rapid temporal changes. Using transitional click trains with subtle inter-click-interval (ICI) shifts, we first show that increasing echo strength systematically enlarges the sound-level step at ICI transitions in both recorded and simulated stimuli, sharpening the physical boundary between segments of distinct temporal structure. In humans, psychophysical detection of ICI changes improves monotonically with echo level in both simulated and real free-field environments, while a late EEG change response scales with echo strength as onset responses remain comparatively stable. In awake rats, electrocorticography over auditory cortex reveals parallel echo-dependent enhancement of transition-evoked activity. Single units exhibit echo-level–dependent amplification of change-related firing and improved neurometric discriminability. Laminar local field potential and current source density analyses further show that echo-dependent divergence emerges first in granular and infragranular layers before propagating to supragranular cortex, consistent with thalamocortical drive followed by intracortical amplification. Together, these findings establish EFTS as a cross-species mechanism that reframes echoes as structured signals the brain uses to sharpen temporal integration in everyday listening.

## Introduction

The auditory system is fundamentally attuned to the temporal structure of sound, which serves as a cornerstone of auditory cognition and enables the perception of complex acoustic phenomena such as speech, music, and environmental signals [1–3]. Temporal structure—encompassing the precise timing, duration, and sequencing of acoustic events—underpins essential cognitive processes, including the segmentation of speech into phonemes and syllables, the discrimination of rhythmic patterns in music, and the localization of transient sounds in noisy environments [4–6]. Without robust sensitivity to temporal cues, the brain would struggle to form coherent auditory objects, leading to fragmented perceptions that impair communication, navigation, and social interaction [7, 8]. In particular, the auditory cortex is adept at extracting behaviorally relevant features from complex acoustic streams, a process crucial for communication behaviors as demonstrated in models of vocal perception [9, 10]. Indeed, deficits in processing temporal structure are implicated in a range of neurodevelopmental and neurological conditions, such as dyslexia, where impaired phonemic timing contributes to reading difficulties, and central auditory processing disorder, which disrupts the interpretation of rapid sound sequences [11, 12]. These impairments highlight the critical role of temporal structure not only in basic sensory encoding but also in higher-order cognitive functions, including attention, memory, and language acquisition [13, 14]. While frequency-based processing has been extensively mapped, how the brain maintains temporal precision in realistic acoustic environments remains a central and unresolved question [15].

Echoing, or reverberation—the persistence of sound due to reflections from surrounding surfaces—is ubiquitous across natural and built environments, from forests and caves to rooms and urban streets [16, 17]. Virtually all auditory experiences occur in reverberant spaces, where direct sound waves intermingle with delayed replicas, altering the temporal envelope and spectral content [18]. Despite its pervasiveness, the functional role of echoing in auditory cognition remains poorly understood. Traditional perspectives frame reverberation as a perceptual challenge, positing that it degrades temporal resolution by smearing fine details and introducing masking noise [19]. Yet, emerging evidence suggests potential adaptive benefits: moderate echoing may enhance signal redundancy, filling temporal voids and bolstering perceptual stability in noisy contexts [20, 21]. This tension reveals a deeper conceptual gap: does the auditory system merely tolerate reverberation, or does it exploit echoes as part of a general computational principle for dealing with natural temporal variability? Surprisingly, whether reverberation can systematically enhance sensitivity to rapid temporal structure—and whether such enhancement reflects a dedicated neural computation—remains unknown, despite its relevance for nearly all real-world listening.

Here we show that reverberation plays a previously unrecognized functional role in auditory processing: it enhances the brain’s sensitivity to rapid temporal transitions. We identify a general phenomenon, echo-facilitated temporal sensitivity (EFTS), in which increasing echo strength selectively amplifies neural and perceptual responses to temporal change while leaving sound-onset encoding comparably stable. Using transitional click trains with subtly different inter-click intervals (ICIs), we find in humans that echoing improves behavioral change detection and EEG change responses, and in rats that electrocorticogram (ECoG) and single-unit activity show echo-level–dependent increases in transition-evoked firing and ROC-based discriminability. Laminar local field potential (LFP) and current source density (CSD) analyses further reveal that echo-dependent divergence emerges first in granular and infragranular layers and then in supragranular layers, consistent with thalamocortical drive followed by intracortical amplification. Together, these findings establish EFTS: reverberation acts as a natural enhancer of temporal integration, sharpening cortical representations of rapid pattern changes in real-world listening.

## Results

### Physical characterization of reverberation-induced level changes

To probe how echoes shape the physical, perceptual, and neural representation of temporal structure, we used transitional click trains (Fig. 1A). Each stimulus comprised two consecutive regular click trains with different inter-click intervals (ICIs). The first segment (e.g., 4-ms ICI, “Reg4”) established a stable temporal pattern, and the second segment (e.g., 4.06-ms ICI, “Reg4.06”) introduced a step change in temporal structure (“Reg_4–4.06_”). This concatenation created a well-defined transition point whose physical consequences could be quantified and then related to behavior and neural activity in the subsequent experiments.

**Figure 1.**
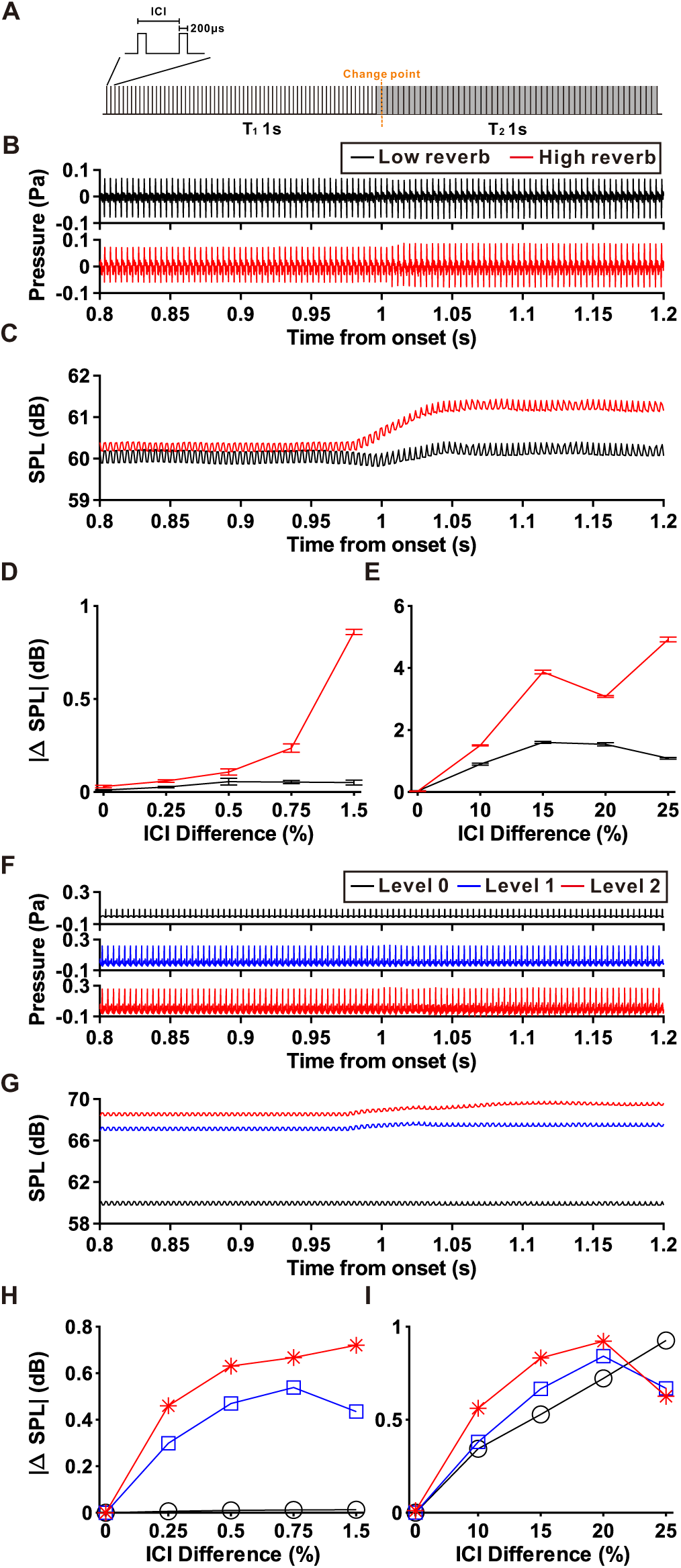
The effect of echoes on the physiological characteristics of transitional click train. **(A)** Layout of the auditory stimulation setup (transitional click train). Two click trains, each comprising 0.2-ms monopolar pulses with different inter-click intervals (ICIs), were linked seamlessly to create a transitional click train. For instance, one at 4 ms—train 1 (T_1_) and the other at 4.06 ms—train 2 (T_2_) were referred to as Reg_4-4.06_. **(B)** Recorded sound pressure of Reg_4-4.06_ used in human Experiment 3 (free-field change detection task, see Fig. 2F) under low-reverb (black) and high-reverb (red) environments. **(C)** Time-varying intensity of the recorded Reg_4-4.06_ computed with a 50-ms bin and a 1-ms step (black: low-reverb; red: high-reverb). **(D)** Absolute intensity difference between pre- and post-change windows (0.9–1.0 s vs. 1.0–1.1 s) as a function of ICI difference in human Experiment 3 (black: low-reverb; red: high-reverb). **(E)** Absolute intensity difference between pre- and post-change windows as a function of ICI difference for the rat ECoG experiment (black: low-reverb; red: high-reverb). **(F)** Simulated sound pressure of Reg_4-4.06_ used in human Experiment 1 (ICI-contrast task, see Fig. 2A) under three simulated echo levels (black: level 0; blue: level 1; red: level 2). **(G)** Time-varying intensity of the simulated Reg_4-4.06_ computed with a 50-ms bin and a 1-ms step (black: level 0; blue: level 1; red: level 2). **(H)** Absolute intensity difference between pre- and post-change windows as a function of ICI difference for human Experiment 1 (black: level 0; blue: level 1; red: level 2)**. (I)** Absolute intensity difference between pre- and post-change windows for the rat ECoG experiment (black: level 0; blue: level 1; red: level 2).

We first characterized how reverberation alters the acoustics of these transitional click trains. For each stimulus, we measured the change in sound level (|ΔSPL|, dB) between short windows immediately before and after the transition. For sounds recorded in real rooms for the human free-field experiments, waveforms from the high-reverb environment showed stronger amplitude modulation than those from the low-reverb condition (Fig. 1B). The corresponding intensity trace exhibited a more pronounced step at the transition under high reverberation (Fig. 1C). Across a range of ICI differences, |ΔSPL| was larger for high than for low reverberation at every contrast (Fig. 1D), indicating that reverberation increases the physical level contrast associated with temporal-structure changes. A second recorded stimulus set, designed for the rat ECoG free-field experiment and spanning larger ICI differences, showed the same pattern: high-reverb recordings consistently produced greater |ΔSPL| than low-reverb recordings (Fig. 1E).

To test whether this effect reflects a general acoustic consequence of echoing rather than idiosyncrasies of a particular room, we next examined simulated reverberation. Transitional click trains were synthesized and then passed through echo filters corresponding to three reflection levels (Levels 0–2), which were later used in the human earphone EEG experiment. As echo level increased, the simulated waveforms showed successively stronger amplitude modulation (Fig. 1F), and the intensity traces displayed progressively larger steps at the transition (Fig. 1G). Across ICI contrasts, |ΔSPL| increased monotonically from Level 0 to Levels 1 and 2 (Fig. 1H). A parallel simulated stimulus set for the rat single-unit experiment yielded qualitatively similar results (Fig. 1I): stronger echoes produced larger level changes around the transition for all but the largest ICI contrast, where the function bent slightly downward, suggesting a modest non-linearity at very large temporal differences.

These analyses focus specifically on changes in overall sound level and do not exclude contributions from other acoustic factors—such as alterations in temporal envelope shape, fine structure, or spectral content—that may covary with echo strength. Nevertheless, across recorded and simulated sounds, in both human and rat stimulus sets, reverberation systematically enhanced the physical intensity change at temporal transitions. This provides a principled acoustic baseline for interpreting the neural and behavioral effects of echo-facilitated temporal sensitivity in the following sections.

### Reverberation improves perceptual detection of temporal changes in humans

We next asked whether reverberation improves perceptual sensitivity to temporal-structure transitions. Human listeners heard transitional click trains in which two segments with slightly different ICIs (e.g., 4→4.01 ms, “Reg_4–4.01_”; 4→4.06 ms, “Reg_4–4.06_”) were concatenated (Fig. 2A). In Experiment 1, participants performed a yes/no change-detection task over insert earphones while we parametrically manipulated simulated reverberation at three reflection levels—little, moderate, and strong (Levels 0–2; Figs. 1F-H and 2B). Change-detection performance increased monotonically with echo level (Fig. 2C; F(2, 20) = 119.05, p < 0.001, two-way repeated-measures ANOVA, Greenhouse–Geisser corrected), indicating that reverberation sharpens perceptual sensitivity to fine temporal variations.

**Figure 2.**
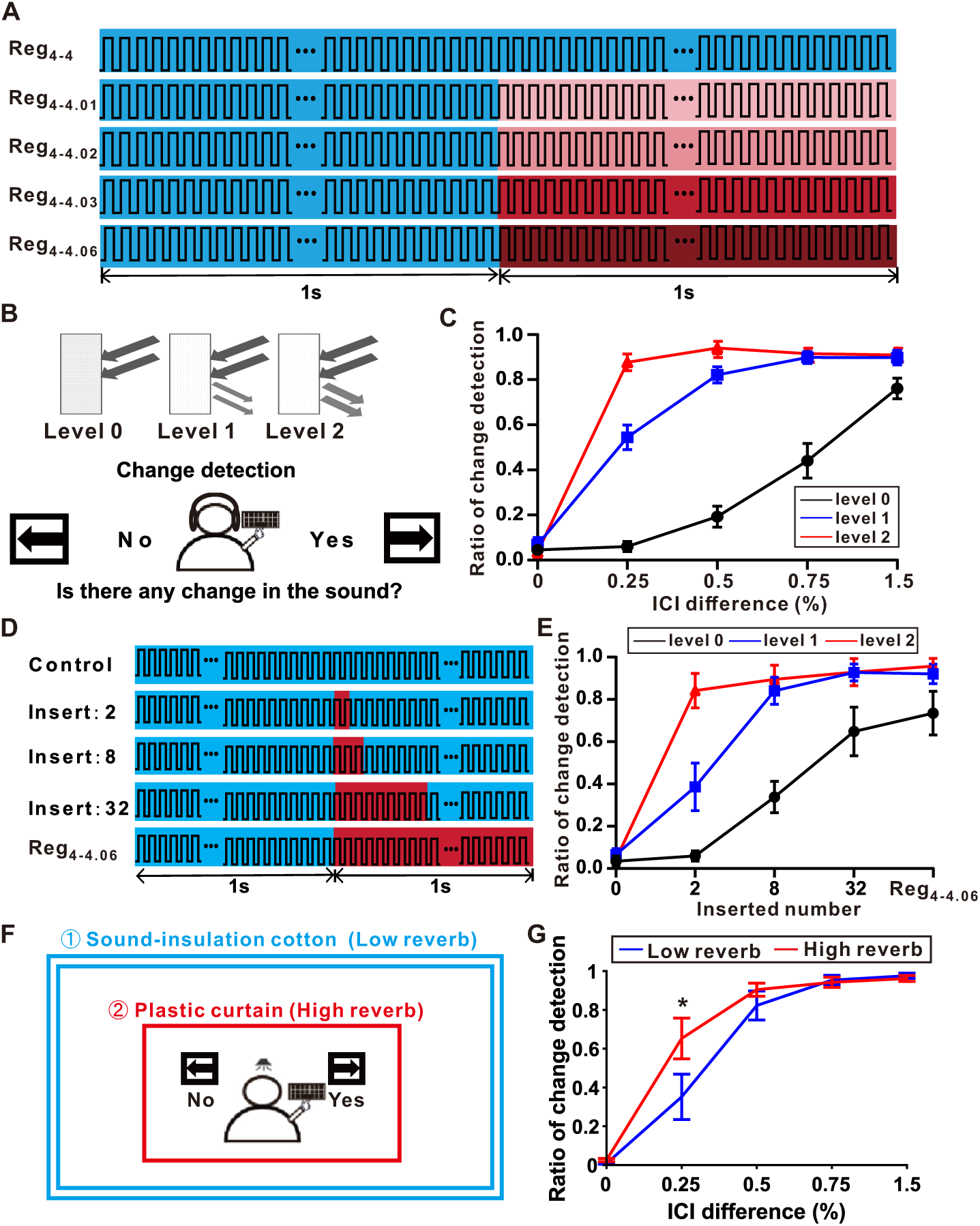
Experimental setup and psychological results of three change detection tasks. **(A)** Schematic diagram of the setup of Experiment 1 (Interval contrast). Five stimuli (Reg_4-4_, Reg_4-4.01_, Reg_4-4.02_, Reg_4-4.03_, and Reg_4-4.06_) were presented in a random order. **(B)** Participants were instructed to indicate whether a change occurred across the transitional click train via button presses. Two simulated echo conditions (level 1 and level 2) and one control condition (level 0) were played through an earphone. **(C)** Ratio of change detection as a function of the ICI differences under level 0 (black), level 1(blue) and level 2 (red) conditions (n = 11), error bars indicate standard error. **(D)** Schematic diagram of the setup of Experiment 2 (Interval contrast). Four distinct interval numbers (0, 2, 8 and 32 intervals of 4.06 ms) were inserted at the 1-second mark within a 2-second regular click train maintaining a constant ICI of 4 ms (shown in the top four rows). For control, the top row illustrates Reg_4–4.06_, composed of a 1-second click train with 4-ms ICI, followed by a 1-second train with 4.06-ms ICI. These five patterns were presented in a randomized order. **(E)** Ratio of change detection as a function of the inserted number under level 0 (black), level 1(blue) and level 2 (red) conditions (n = 13), error bars indicate standard error. **(F)** Schematic diagram of the setup of Experiment 3 (Open field). Participants listen to stimulation from a speaker in front of them. Two experimental environments with different echo characteristics were established by using materials with distinct acoustic properties. In one condition, the experiment was conducted in a soundproof room covered with sound-insulation cotton, which provided a low-reverb environment (blue). In the other condition, the participant was surrounded by a circle of plastic curtains, forming a high-reverb environment (red). The sounds were the transitional click trains with the same ICI contrast setup as Experiment 1 without any modification. **(G)** Ratio of change detection as a function of the inserted number under low-reverb (blue) environment and high-reverb (red) environment (n = 10), error bars indicate standard error.

To probe how echoes influence temporal integration, Experiment 2 inserted 0 (control), 2, 8, or 32 intervals of 4.06 ms into an otherwise regular 4-ms train and compared these conditions with the full Reg_4–4.06_ transition at each echo level (Fig. 2D). Behavioral performance improved systematically with both insertion number and reverberation strength (Fig. 2E; F(2, 24) = 20.37, p < 0.001 for echo strength; F(4, 48) = 46.74, p < 0.001 for insertion number; two-way repeated-measures ANOVA, Greenhouse–Geisser corrected). Under little echoing, short insertions (2 and 8 intervals) were rarely detected, whereas under strong echoing they were reliably perceived, suggesting that reverberation facilitates the accumulation of brief deviations into a coherent percept of change.

Finally, to assess ecological generality, Experiment 3 repeated the change-detection task in real free-field environments. Participants listened either in a soundproof room lined with sound-insulation cotton (low-reverb condition) or within a reflective chamber formed by plastic curtains (high-reverb condition; Figs. 1B-D and 2F). In the reflective chamber, change-detection accuracy at the most challenging contrast (Reg_4–4.01_) was significantly higher than in the low-reverb room (Fig. 2G; p < 0.01, Wilcoxon signed-rank test).Across simulated and real environments, reverberation consistently enhanced behavioral sensitivity to subtle temporal transitions, establishing a robust perceptual manifestation of the echo-facilitated temporal sensitivity (EFTS) revealed by the physical analyses in Figure 1.

### Reverberation Selectively Amplifies Cortical Change Responses in Humans

We then asked whether this perceptual EFTS is mirrored in cortical dynamics, using EEG. In Experiment 1, grand-averaged responses at electrode P4 showed that the onset response (0–300 ms) was largely invariant across echo levels (Fig. 3A; Supplementary Fig. 1A, B), confirming that reverberation did not substantially alter baseline auditory encoding. By contrast, the late change response—previously identified as a neural indicator of temporal integration [22–24]—increased systematically with echo strength (Fig. 3A, >1000 ms). Scalp topographies revealed broader and stronger change-response distributions at higher echo levels, with peaks over temporo-parietal electrodes and a clear dependence on ICI contrast (Fig. 3B).

**Figure 3.**
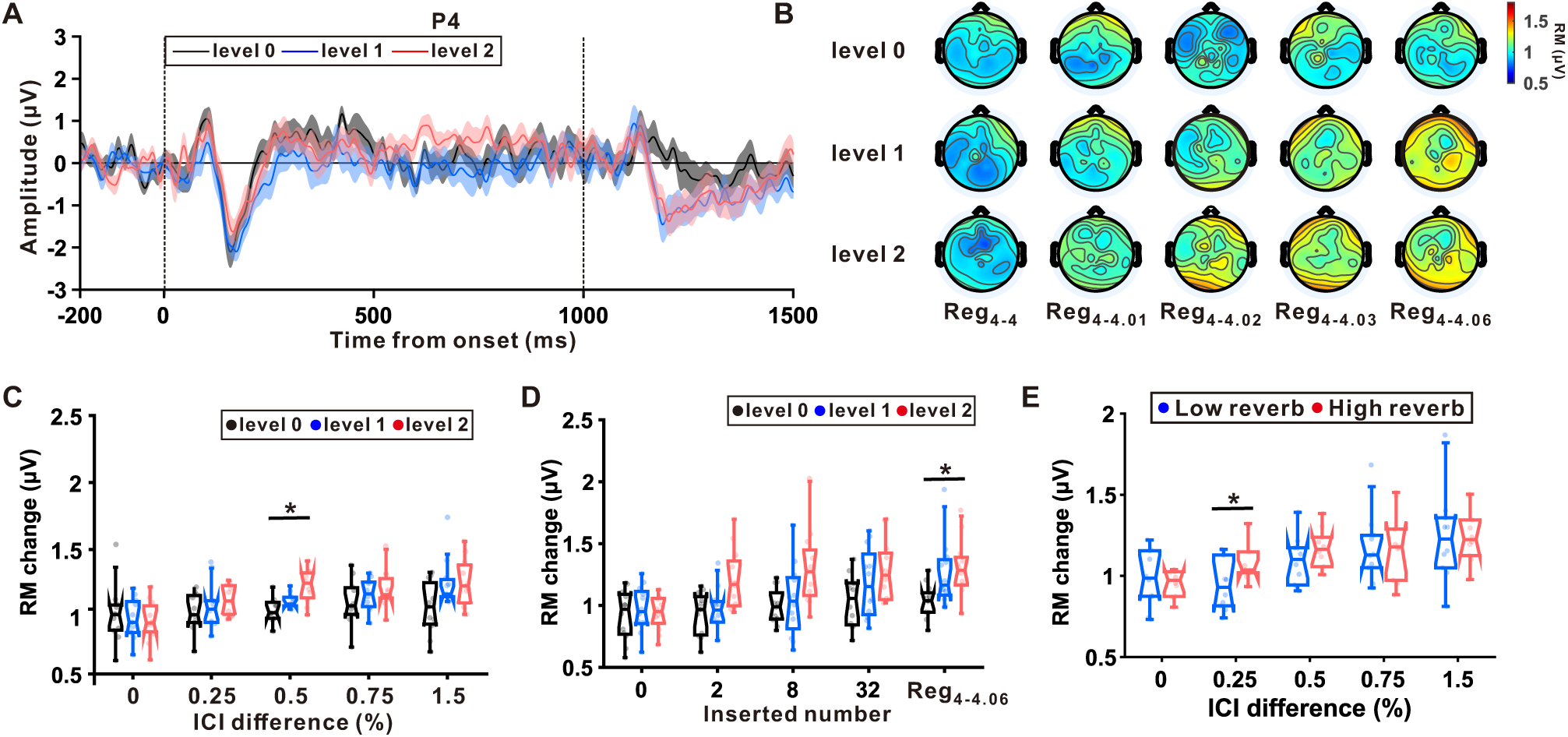
The EEG result of three change detection tasks. **(A)** Grand-averaged responses at electrode P4 across 11 subjects participating in the Experiment 1 (experimental setup is shown in Fig. 2A) under level 0 (black), level 1(blue) and level 2 (red) conditions at Reg_4-4.06._ The shaded area around the curve represents ± SEM. Vertical black dashed lines represent the onset (0) and change (1000) of the transitional click train, respectively. **(B)** Topographic maps show change RM distributions under three echo conditions across the scalp across different ICI contrasts. **(C)** Boxplots show group-level RM values (averaged across electrodes) under level 0 (black), level 1(blue) and level 2 (red) conditions (mean: central line; standard error: box edges; 10^th^-90^th^ percentiles: whiskers). Transparent dots represent individual participants. Asterisks (*) indicate significant differences between groups (p < 0.05, Friedman’s test with multiple comparison). **(D)** Boxplots show group-level RM values (averaged across electrodes) in Experiment 2 (experimental setup is shown in Fig. 2D) under level 0 (black), level 1(blue) and level 2 (red) conditions (mean: central line; standard error: box edges; 10^th^-90^th^ percentiles: whiskers). Transparent dots represent individual participants. Asterisks (*) indicate significant differences between groups (p < 0.05, Friedman’s test with multiple comparison). **(E)** Boxplots show group-level RM values (averaged across electrodes) in Experiment 3 (experimental setup is shown in Fig. 2F) under low-reverb (blue) environment and high-reverb (red) environment (mean: central line; standard error: box edges; 10^th^-90^th^ percentiles: whiskers). Transparent dots represent individual participants. Asterisks (*) indicate significant differences between groups (p < 0.05, paired t-test).

Quantitatively, response magnitudes (RM) of the change component exhibited significant main effects of echo level (Fig. 3C; F(2, 20) = 5.61, p < 0.05, two-way repeated-measures ANOVA, Greenhouse–Geisser corrected) and ICI contrast (F(4, 40) = 8.18, p < 0.001). In Experiment 2, change-response amplitudes scaled with both insertion number and echo strength (Fig. 3D; F(2, 24) = 7.63, p < 0.01 for echo level; F(4, 48) = 8.55, p < 0.001 for insertion number). Across echo levels, the change response increased with insertion number and approached an asymptote near the full Reg_4–4.06_ transition, reinforcing that this component reflects temporal integration of emerging pattern deviations [22,23]. In the free-field experiment (Experiment 3), change-response magnitudes were larger in the high-reverb chamber than in the low-reverb room at the most difficult contrast (Reg_4–4.01_; Fig. 3E; p < 0.05, Wilcoxon signed-rank test), whereas onset responses remained comparable across environments (Supplementary Fig. 1C, D; p = 0.91, Friedman’s test; p = 0.10, Wilcoxon signed-rank test).

Together, these EEG results show that reverberation selectively boosts cortical responses to temporal-structure transitions while leaving onset encoding relatively stable. In conjunction with the behavioral data, they demonstrate a human expression of EFTS, in which echoes act as an environmental resource that the auditory system uses to enhance neural and perceptual sensitivity to rapid temporal changes.

### Cross-species Conservation of EFTS in Rat Auditory Cortex

Building on the human findings, we next examined whether EFTS is conserved across species. We recorded ECoG signal from awake rats exposed to transitional click trains in free field under two acoustic environments: a soundproof room (low-reverb condition) and, for comparison, a reflective chamber constructed from a corrugated cardboard box mounted on the experimental platform to produce natural reverberation (high-reverb condition) (Fig. 4A). The transitional sequences concatenated segments with ICIs of 4→4.4–5 ms (Reg_4–4_, Reg_4–4.4_, Reg_4–4.6_, Reg_4–4.8_, Reg_4–5_; Fig. 4B).

**Figure 4.**
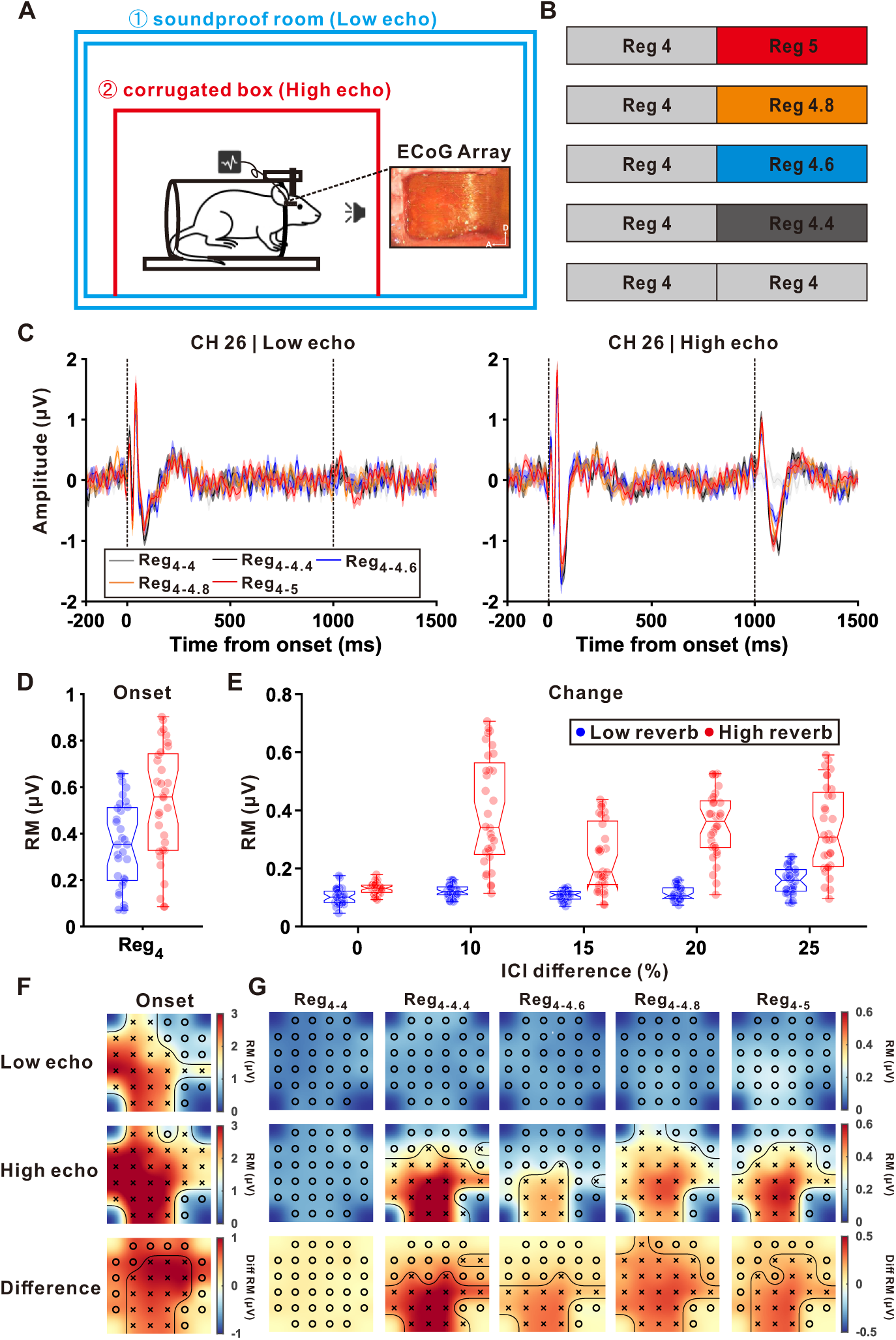
ECoG results from an example rat in two echo environments. **(A)** Schematic diagram of the experimental setup. The rat was head-fixed facing a speaker. The two echoic environments were configured differently. A soundproof room lined with sound-insulation cottom was used to create a low-reverb environment, while a corrugated cardboard box mounted on the experimental platform was used to construct a high-reverb environment. A photo of rat auditory cortex (AC) covered with ECoG array was displayed. **(B)** Transitional click trains used in the experiment including five ICI contrasts: Reg_4-4_, Reg_4-4.4_, Reg_4-4.6_, Reg_4-4.8_, and Reg_4-5_. The five sounds were presented in a random order. **(C)** Left: Averaged responses at channel 26 across repeated trials under low-reverb environment. Five curves corresponding to different ICI contrasts were colored differently: Reg_4-4_ (gray), Reg_4-4.4_ (black), Reg_4-4.6_ (blue), Reg_4-4.8_ (orange), and Reg_4-5_ (red). The shaded area around the curve represents ± SEM. Vertical black dashed lines represent the onset (0) and change (1000) of the transitional click train, respectively. Right: Averaged responses at channel 26 across repeated trials under high-reverb environment. **(D)** Boxplots show RM values of onset responses under low-reverb (blue) environment and high-reverb (red) environment (mean: central line; standard error: box edges; 10^th^-90^th^ percentiles: whiskers). **(E)** Boxplots show RM values of change responses under low-reverb (blue) environment and high-reverb (red) environment (mean: central line; standard error: box edges; 10^th^-90^th^ percentiles: whiskers) across five ICI differences. **(F)** Topographic maps show the distribution of onset RM under low-reverb environment (top) and high-reverb environment (middle) and ΔRM (bottom). Channels with significant RM and ΔRM were highlighted with ‘x’ (p < 0.05, nonparametric bootstrap procedure). **(G)** Topographic maps show the distribution of change RM under low-reverb environment (top) and high-reverb environment (middle) and ΔRM (bottom) across different ICI contrasts. Channels with significant RM and ΔRM were highlighted with ‘x’ (p < 0.05, nonparametric bootstrap procedure).

In the soundproof room, ECoG recordings revealed a robust onset response but minimal change-related activity in a representative channel (Fig. 4C, left). By contrast, in the reflective chamber, both onset and change responses were clearly expressed (Fig. 4C, right). Quantitative analysis confirmed a larger onset amplitude in the reflective condition (Fig. 4D; p < 0.001, paired t-test). More importantly, change-response amplitudes increased markedly under the reflective condition across all non-zero ICI contrasts (Fig. 4E; p < 0.001 for all conditions). Thus, mirroring the human results, reverberation selectively enhances cortical sensitivity to temporal pattern changes in rats, consistent with EFTS.

Spatial mapping of ECoG activity further illustrated EFTS across the cortical surface. Topological distributions of onset responses were similar across environments but showed higher overall amplitudes in the reflective chamber (Fig. 4F). In contrast, change-response topographies differed strikingly: whereas the low-reverb condition produced weak, spatially confined activation, strong echoing yielded broader and stronger cortical responses (Fig. 4G). Difference maps (bottom rows of Fig. 4F and G, p < 0.05, nonparametric bootstrap procedure with 1,000 iterations) highlighted clear reverberation-driven enhancement over auditory cortex. The EFTS effect on the change response was also consistently observed in another rat (Supplementary Fig. 2). In addition, a supplementary ECoG experiment using simulated sounds delivered through closed-field earphones further verified the robustness of the echo enhancement (Supplementary Fig. 3; see Methods for details).

Together, these findings demonstrate that EFTS is conserved across species: reverberation strengthens change responses in rat auditory cortex, paralleling the echo-facilitated enhancement of temporal processing observed in humans.

### EFTS at the neuronal level in rat auditory cortex

Building on the ECoG findings, we recorded single-unit activity in rat AC during transitional click trains delivered via closed-field earphones while parametrically manipulating simulated reverberation (little, moderate, strong; Levels 0–2), identical to the human settings. We reasoned that if EFTS indeed reflects a sharpening of cortical temporal processing, it should be evident at the level of single neurons, which are known to exhibit robust plasticity in their representation of learned or behaviorally relevant complex sounds [25]. For a representative AC neuron, raster plots for Reg_4–4.8_ showed progressively stronger firing following the temporal structure transition as echo level increased (Fig. 5A, top). The corresponding peristimulus time histograms (PSTHs) confirmed this pattern, with a clear enhancement of the change response under stronger echoing (Fig. 5A, bottom). For this neuron, change-evoked firing rates scaled monotonically with echo level across all tested contrasts, demonstrating that greater reverberation consistently amplifies neural sensitivity to temporal change (Fig. 5B; F(2,433) = 30.03, p < 0.001 for echo strength; F(4, 433) = 8.25, p < 0.001 for ICI contrast; two-way ANOVA).

**Figure 5.**
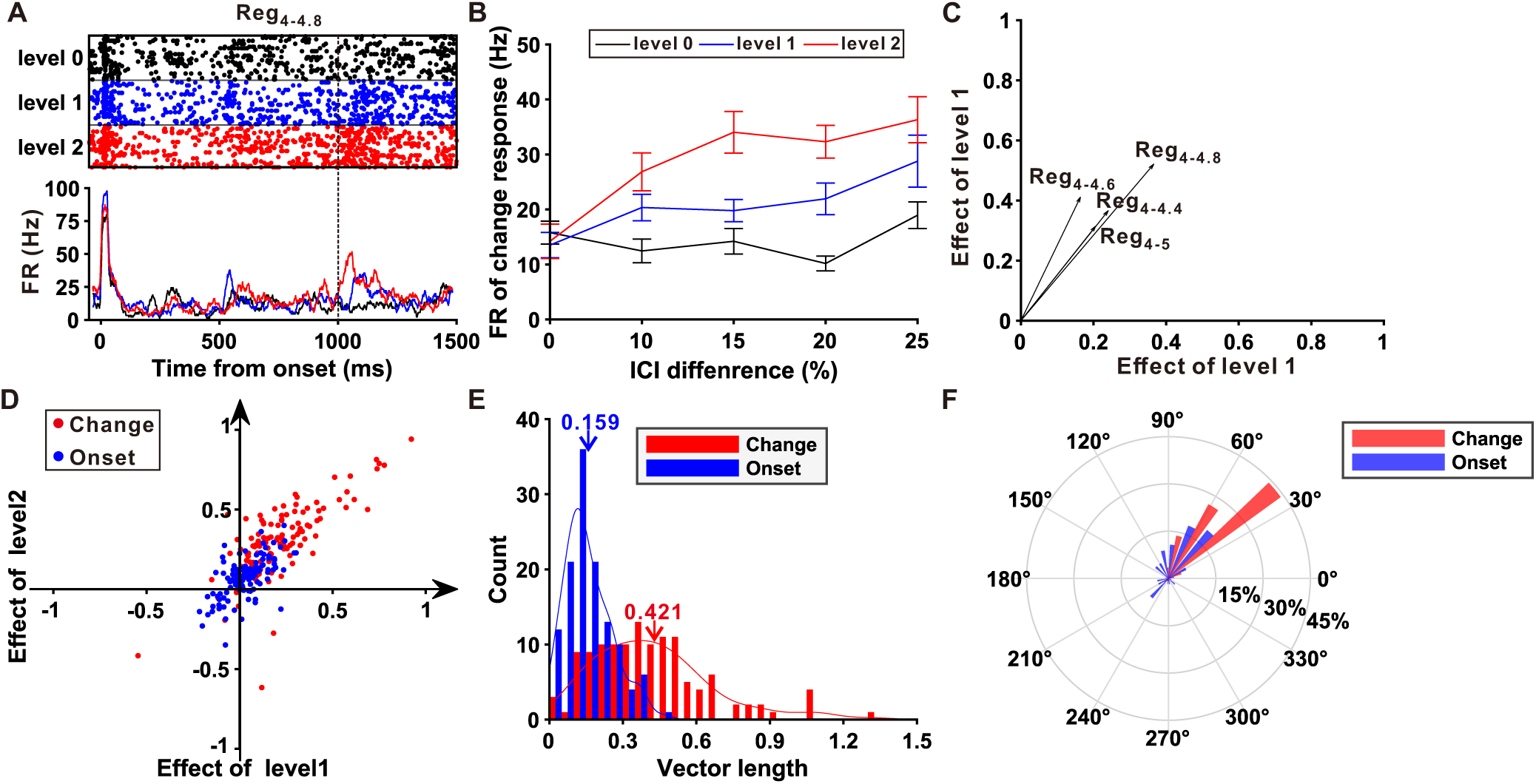
The effect of echo on neuronal responses. **(A)** Top: Raster plots illustrate neuronal responses to Reg_4-4.8_ under three echo levels from an example AC neuron. Bottom: Peri-stimulus-time histograms (PSTHs) show the response pattern of the example neuron across the sound. The zero point corresponds to the onset of the sound. **(B)** Firing rate of change response as a function of the ICI difference under level 0 (black), level 1(blue) and level 2 (red) conditions, error bars indicate standard error. **(C)** The effect vectors of four ICI contrasts. **(D)** The distribution of population (n = 124) effect vectors of onset responses (blue) and change responses (red) under the ICI contrast of Reg_4-4.8_. **(E)** The distribution of vector length for onset responses (blue) and change responses (red) under the ICI contrast of Reg_4-4.6._ **(F)** The distribution of vector phases in a polar coordinate for onset responses (blue) and change responses (red) under the ICI contrast of Reg_4-4.6_.

To quantify modulation at the single-unit level, we defined an effect index for each echo condition as the normalized difference between the change response at Level 1 (or Level 2) and the response at Level 0 (control). Plotting the Level-2 effect against the Level-1 effect yielded a vector whose length represented the magnitude of EFTS and whose angle indicated the modulation direction (amplification vs. suppression). For the example neuron at Reg_4–4.8_ ms, the vector lay in the first quadrant with substantial magnitude, indicating robust EFTS-driven amplification of the change response (Fig. 5C).

At the population level for Reg_4–4.8_, vectors for change responses lay predominantly in the first quadrant and farther from the origin than vectors for onset responses (Fig. 5D). Consistently, histogram comparisons showed significantly greater vector lengths for change than for onset responses (Fig. 5E; p < 0.001, paired t-test), indicating stronger EFTS for transition-related activity. Vector-angle distributions also differed systematically between components: change responses clustered at amplification angles, whereas onset responses exhibited a more uniform spread across angles (Fig. 5F; p < 0.001, paired circular test, Rayleigh test on angular differences). Population analyses at other contrast conditions yielded comparable results (Supplementary Fig. 4). Collectively, these findings demonstrate that reverberation enhances firing related to temporal-structure transitions at the single-unit level, providing direct evidence that EFTS selectively amplifies neural representations of temporal change in the AC.

### Functional significance of EFTS

To assess the functional consequence of EFTS, we tested whether reverberation improves neural discriminability of temporal change using Receiver Operating Characteristic (ROC) analysis. For each echo level (little, moderate, strong; Levels 0–2), we compared spike-count distributions from the change period with those from the preceding control period for Reg_4-4.8_ (Fig. 6A). The separation between distributions increased systematically with echo level, indicating improved discrimination under stronger reverberation.

**Figure 6.**
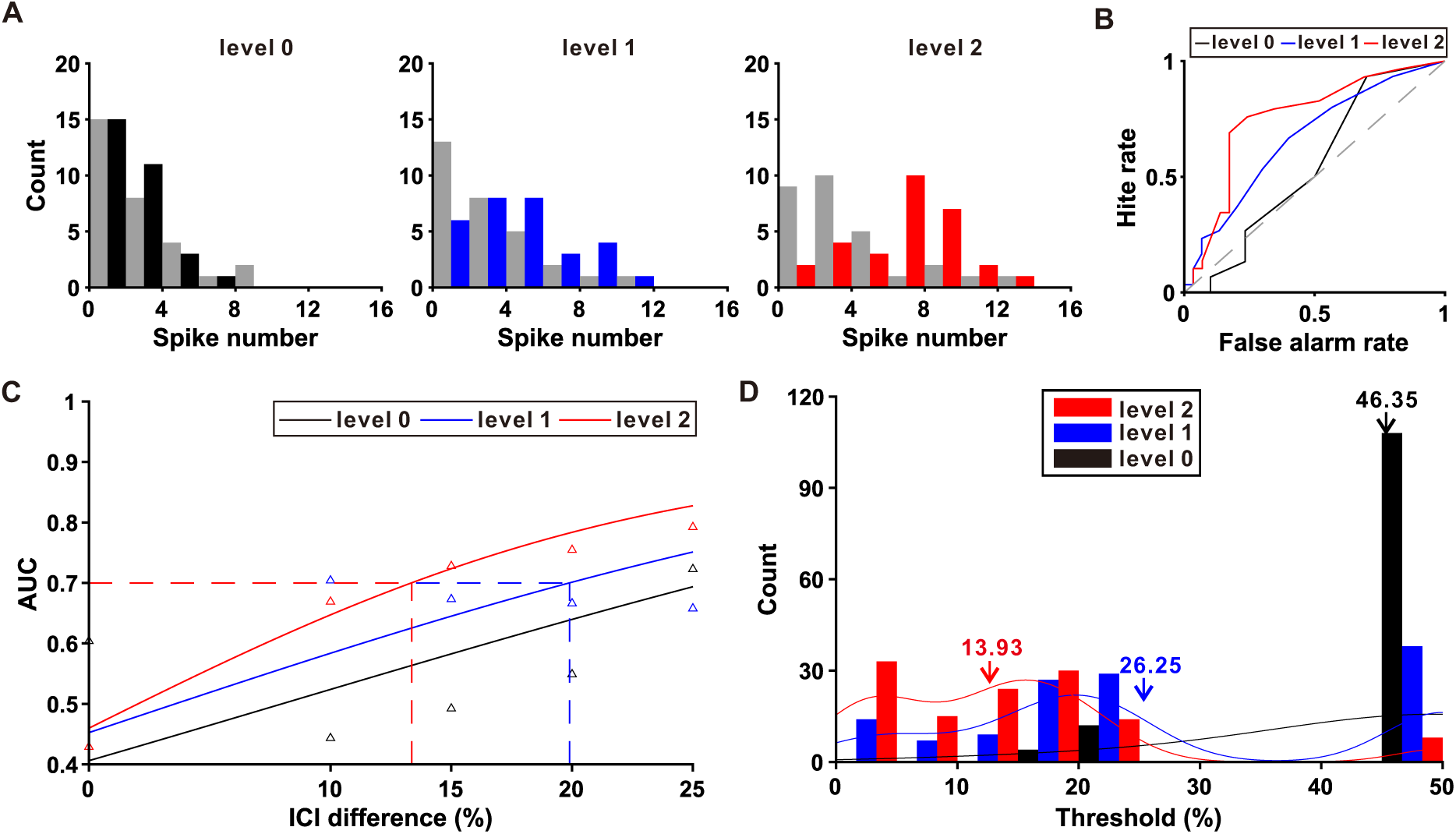
Neuronal change detection threshold decreases with the increase of echo level. **(A)** The comparisons between the distribution of neuronal responses during the response window (0 to 200 ms) and baseline window (−200 to 0 ms) for three echo conditions under Reg_4-4.8_ from the same example neuron as Figure 3. Red: response; blue: baseline. **(B)** ROC analysis of comparison in A between two choices across three echo conditions. Black: level 0, blue: level 1, red: level 2. **(C)** Neurometric function showing the proportion of change detection of an ideal observer as a function of the ICI contrasts. Each data point corresponds to an AUC value computed from a pair of firing rate distributions obtained from the response window and baseline window, like those shown in each column in B. Three lines show the cumulative Gaussian fits under three echo conditions: level 0 (black), level 1 (blue) and level 2 (red). The dashed lines show the neuronal thresholds that reaches 0.7. **(D)** Distribution of thresholds under three echo levels: level 0 (black), level 1 (blue) and level 2 (red).

Consistent with this, ROC curves showed steeper hit-rate growth and reduced false-alarm rates as echo increased (Fig. 6B), and the area under the curve (AUC) rose monotonically with echo level. Extending across ICI contrasts, we computed AUCs to construct neurometric functions and defined the discrimination threshold as the ICI difference yielding 70% correct (Fig. 6C). In the example neuron, higher echo produced steeper neurometrics and smaller thresholds, reflecting enhanced neural sensitivity to temporal change.

At the population level, thresholds shifted lower with increasing echo: means decreased from 46.35% (Level 0) to 26.25% (Level 1) and 13.93% (Level 2). A Friedman’s test confirmed a significant main effect of Echo Level (Fig. 6D; p < 0.001, Friedman’s test). Together, these results show that EFTS not only amplify change-evoked responses but also improves the fidelity of cortical encoding for temporal differences in reverberant environments.

### Neuronal mechanism of EFTS across cortical layers

To uncover the cortical circuitry underlying EFTS, we analyzed laminar LFPs recorded with linear probes spanning all cortical layers. An example response to a transitional click train (Reg_4-4.8_, Level 2) shows distinct temporal profiles across depth (Fig. 7A). CSD analysis (Fig. 7B) identified three zones: supragranular (Sg), granular (Gr), and infragranular (Ig) layers.

**Figure 7.**
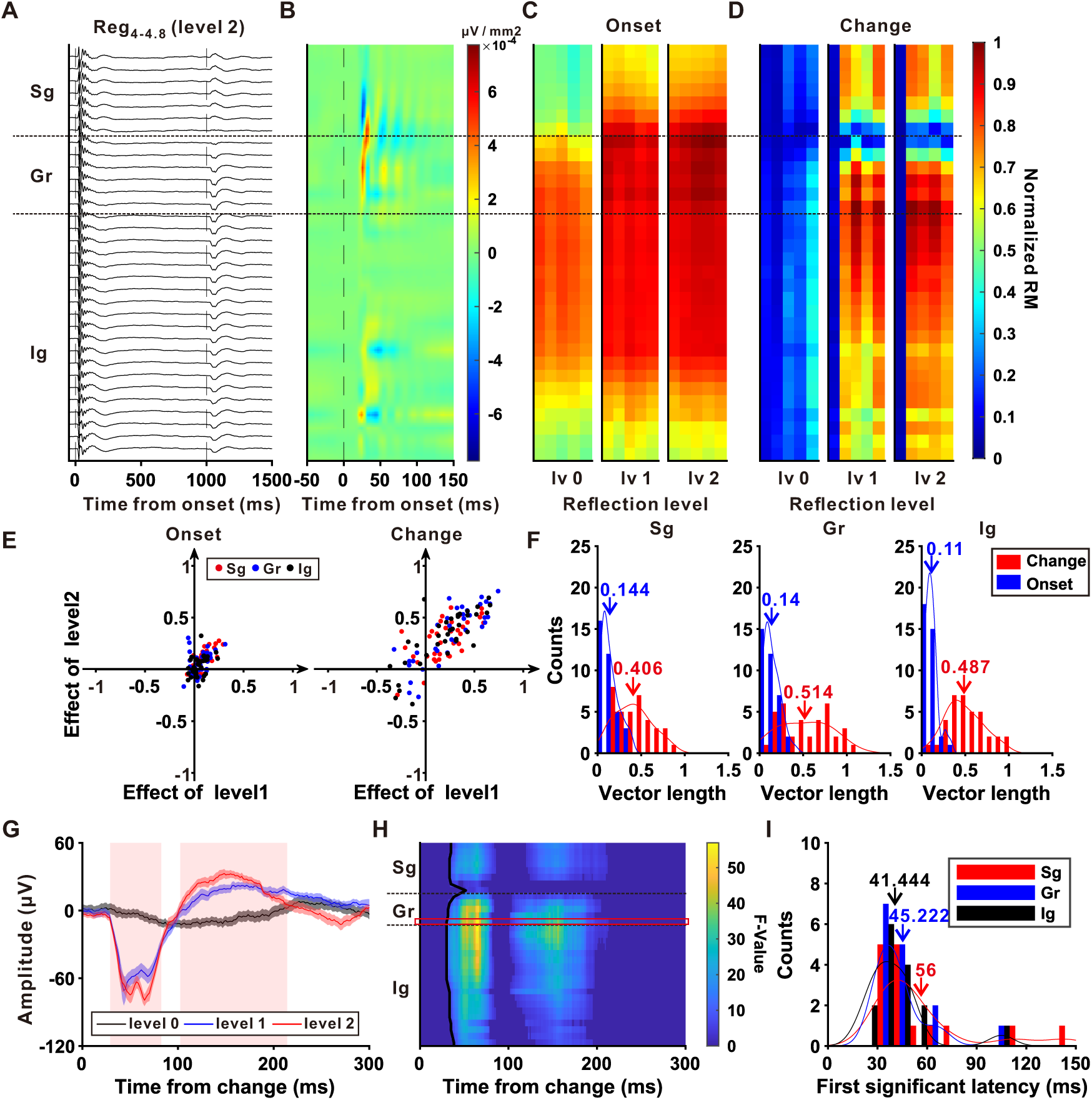
Laminar profiles of echo effect on onset and change responses in the auditory cortex. **(A)** LFP responses to Reg_4-4.4_ throughout different layers of an example penetration in auditory cortex. Vertical dashed lines marked onset of the sound and change point separately. **(B)** Current source density (CSD) map computed from the LFP of the example penetration. **(C)** The distribution of normalized onset RM across cortical layers. Each of the three subplots represents a distinct echo condition, and within each subplot, the five columns correspond to different ICI contrasts. **(D)** The distribution of normalized change RM across cortical layers. **(E)** The distribution of effect vectors of Reg_4-4.8_ throughout different layers (red: Sg; blue: Gr; black: Ig) for onset (left) and change (right) responses, based on 36 penetrations in the AC. **(F)** The distribution of vector lengths for onset (blue) and change (red) responses across layers Sg (left), Gr (middle) and Ig (right). Arrows denote mean values. **(G)** The average LFP change response to Reg_4-4.8_ from an example channel in Gr (marked with a red rectangle in H) for level 0 (black), level 1 (blue) and level 2 (red). The shaded area around the curve represents ± SEM. **(H)** F-values of response comparisons among three echo levels for channels in the example channel from AC, aligned to the change point. Colors other than dark blue indicate significant time bins (permutation test, p < 0.05, FDR corrected for channels and time bins). The black curves indicate the boundaries of the first significant time bins. **(I)** Histogram of minimum time bins when the three traces in G start to show significant differences for different layers (black: Ig, blue: Gr, red: Sg) in penetrations where all three layers exhibit significant echo effect. The Gr showed lowest echo effect. (Friedman’s test with multiple comparison, p < 0.01).

For the onset response, activity was predominantly localized to Gr and Ig under little echoing and increased modestly across echo levels (little, moderate, strong; Fig. 7C). By contrast, the change response exhibited clear echo dependence: under little echoing, change-related activation was weak, whereas with increasing reverberation the response amplitude rose markedly across all three layers (Fig. 7D). The average response across all penetrations is shown in Supplementary Fig. 5. These observations suggest that EFTS engage distributed laminar processing that amplifies transition-related activity throughout the cortical column.

Quantitative analyses corroborated this pattern. Plotting Level-2 versus Level-1 effects for each layer (Reg_4-4.8_) showed onset-response vectors clustered near the origin, indicating minimal echo modulation (Fig. 7E, left), whereas change-response vectors displayed larger magnitudes across layers (Fig. 7E, right). Distributions of vector lengths revealed significantly stronger EFTS for change than for onset responses within all layers (Fig. 7F, p < 0.001, paired t-test), and other ICI contrasts yielded comparable results (Supplementary Fig. 6).

We next examined temporal dynamics by comparing F-values across echo levels for the change response to Reg_4-4.8_ to estimate the earliest significant time for each channel (permutation-based ANOVA with 1000 iterations, p < 0.05). A representative channel illustrates that enhancement under stronger echoing emerged rapidly after the transition onset (Fig. 7G). The earliest significant divergence occurred in Gr and Ig (black trace, Fig. 7H), consistent with an initial thalamocortical input. Population latencies for first significance confirmed that Sg lagged behind Gr and Ig (Fig. 7I, means: Gr 45.22 ms, Sg 56 ms, Ig 41.44 ms, p < 0.01, Friedman’s test with multiple comparison), but there was no difference between Gr and Ig (p = 0.98, Friedman’s test with multiple comparison). The results of other ICI contrasts are shown in Supplementary Fig. 7. The enhancement effect on neural discriminability across laminar zones showed a similar pattern: Gr exhibited the lowest threshold, whereas Sg was less sensitive to the temporal change in the echoing conditions (Supplementary Fig. 8). Together, these findings indicate that EFTS arises early within the cortical column—likely initiated by thalamocortical input to Gr—and then propagates to superficial layers via intracortical recurrence, recruiting a vertically integrated network that selectively enhances change-related processing.

## Discussion

Our investigation elucidates EFTS, whereby reverberation augments sensitivity to rapid changes in temporal structure. At the physical level, we showed that echoes systematically increase the absolute change in sound level at temporal transitions: for both recorded and simulated transitional click trains, the intensity step between pre-and post-change windows was larger under higher echo levels across the ICI contrasts and stimulus sets used for humans and rats, with only a slight non-linearity at the largest contrast (Fig. 1). In humans, psychophysical detection of subtle ICI transitions improved monotonically with echo level in both simulated and real acoustic environments (Fig. 2) and was paralleled by selective amplification of a late EEG change response, while onset encoding remained comparatively stable (Fig. 3). This facilitation was conserved in awake rats, where ECoG over auditory cortex showed analogous echo-dependent enhancement of change responses and expanded cortical activation in reflective environments (Fig. 4). At the neuronal level, single units in rat auditory cortex exhibited robust echo-driven amplification of transition-related firing (Fig. 5), ROC neurometrics revealed improved discriminability with lower thresholds under stronger reverberation (Fig. 6), and laminar analyses indicated that echo-dependent divergence emerges first in granular and infragranular layers before propagating to supragranular layers (Fig. 7). Together, these results reposition reverberation as an adaptive mechanism that selectively strengthens neural representations of rapid temporal change while preserving relatively stable onset encoding.

### EFTS in the auditory system and cross-species conservation

At the most basic level, EFTS is grounded in a systematic physical effect of reverberation on temporal transitions. Across both recorded and simulated transitional click trains, increasing echo strength consistently enlarged the change in sound level at the ICI transition: intensity steps between pre- and post-change windows were larger under high than low echo for the human and rat stimulus sets, with only a slight non-linearity at the very largest ICI contrast (Fig. 1). These results indicate that reverberation sharpens the acoustic contrast between segments of distinct temporal structure, providing a natural “gain field” at temporal boundaries on which the auditory system can operate. These analyses focus specifically on changes in overall sound level and do not exclude contributions from other acoustic factors—such as alterations in temporal envelope shape, fine structure, or spectral content—that may covary with echo strength.

On this physical foundation, EFTS manifests as a dose-dependent improvement in temporal change detection, likely stemming from reverberation’s provision of redundant temporal cues that reinforce neural adaptation and perceptual coherence [22, 23]. This process aligns with models of auditory scene analysis where the brain constructs a perceptual representation by extracting the statistical structure of “sound textures” over long time scales, upon which discrete temporal “events” are superimposed [26]. The echoes in our stimuli may serve to define and stabilize this statistical background, thereby facilitating the detection of deviations embedded within it. In humans, behavior and EEG scaled monotonically with echo level: detection accuracy rose across little, moderate, and strong echoing (Figs. 2C and 3A-B), while the change response increased without concomitant alterations in the onset response (Fig. 3C and Supplementary Fig. 1B). Insertion manipulations showed that brief substitutions—only 2 or 8 intervals—were sufficient to elicit robust change responses when echoes were present (Fig. 3D), indicating that reverberation promotes the accumulation of fleeting deviations into coherent percepts. Free-field validation confirmed ecological generality: in a reflective chamber, human change detection and change-evoked EEG responses were enhanced at the most challenging contrast (Reg_4-4.01_; Figs. 2G and 3E). These observations support models in which echoes “fill in” temporal gaps and stabilize integrated representations without inflating baseline encoding [18, 20], align with psychophysical evidence that moderate reverberation can bolster speech intelligibility and perceptual stability [27, 28], and are consistent with classic room-acoustics formulations of temporal-envelope transfer [29]. Because listeners can detect subtle shifts in the balance between direct and reflected sound energy [17, 30], this sensitivity may explain how graded changes in reverberation produce the distinct perceptual and neural effects seen across echo levels.

Cross-species conservation is likewise evident in awake rats, where ECoG over AC revealed larger transition-evoked responses in a reflective chamber than in a sound proof room (Fig. 4C–E), with spatial maps highlighting enhancement distribution over auditory cortex (Fig. 4F–G). In line with the human EEG results, this correspondence indicates a phylogenetically preserved cortical mechanism through which reverberation strengthens temporal integration and enhances sensitivity to rapid pattern changes. Consistent with auditory scene analysis, the results suggest that reflections act as structured cues exploited to maintain perceptual continuity in natural environments [5, 31, 32]. Moreover, the consistency between human and rat cortical measures, together with prior demonstrations of temporal-integration signals across species [22, 23], argues that EFTS reflects shared cortical computations—likely involving recurrent dynamics and rhythmic coordination that bind temporally dispersed events into unified percepts [13, 14]. Such mechanisms may also support the spontaneous extraction of abstract auditory categories, as observed in non-human primates [33]. Beyond theoretical significance, EFTS also offers translational implications for echo-aware hearing-aid design and for rehabilitation of temporal-processing deficits such as dyslexia or central auditory processing disorder [11, 12]. In ecological terms, converting reflections from distortions into cues allows the brain to hear echoes as signals, not noise.

### Neuronal Mechanism Underlying EFTS

Our recordings were confined to auditory cortex, within this structure, the neuronal underpinnings of EFTS reflect selective cortical amplification of transition-related activity. This selective enhancement of responses to temporal changes is conceptually aligned with the auditory cortex’s well-established role in detecting unexpected stimuli or stimulus deviations, a key mechanism for highlighting salient information in the acoustic scene [10, 34, 35]. Single-unit examples showed progressively stronger change-related firing with increasing echoing (Fig. 5A) and monotonic scaling across ICI contrasts (Fig. 5B). A vector analysis contrasting Level-1 and Level-2 echoing quantified this selectivity: change-response vectors predominantly fell in the amplification quadrant and were displaced farther from the origin than onset vectors (Fig. 5C–F), indicating that echoes potentiate transition coding while sparing onset encoding. Functionally, ROC analyses demonstrated greater separation between change and preceding spike-count distributions with increasing echo level, resulting in higher AUC values and lower neurometric thresholds (Fig. 6). Thus, within AC, echoes convert acoustic redundancy into a neural advantage by elevating the signal-to-noise ratio for detecting temporal transitions.

Laminar analyses clarify how this amplification unfolds in cortical microcircuits. LFP/CSD recordings revealed that Echo-dependent divergence emerged last in the supragranular layer, reflecting earlier thalamocortical-driven modulation in deeper layers and its vertical propagation (Fig. 7D, 7G–I). While onset activity remained largely stable and weighted to Gr/Ig across echo levels (Fig. 7C), vector-length comparisons showed significantly stronger echoing effects for change responses within every layer (Fig. 7E–F), consistent with a vertically integrated microcircuit in which thalamocortical input is amplified by intracortical recurrence [36–38]. Although we did not record in subcortical nuclei, this laminar timing pattern aligns with hierarchical and recurrent models of auditory cortical processing in which early feedforward drive is sharpened by local and feedback circuitry to enhance sensitivity to emerging temporal structure [14, 15, 39]. Future work should investigate whether specific inhibitory interneuron subtypes within and across layers, which are known to shape cortical response properties and plasticity [40], are critically involved in gating or amplifying the echo-dependent enhancement of change responses. In sum, our AC-focused data indicate that reverberation enhances temporal processing by engaging thalamocortical inputs and cortical recurrence to broaden temporal integration and selectively boost neural representations of change.

## Methods

### Subjects and Surgical Procedures

#### Electroencephalogram (EEG) Participants

We recruited 31 volunteers (16 males and 15 females; *mean age* = 32.9 years, *SD* = 7) with normal hearing to participate in the EEG experiments. Among them, 11 subjects took part in Experiment 1, 12 in Experiment 2, and 10 in Experiment 3. The experimental procedures complied with ethical guidelines and received approval from the Institutional Review Board (approval number: IRB-20230131-R). All participants provided written informed consent before the experiments. Each auditory stimulus was presented at least 40 times per participant across all experimental sessions.

#### Electrocorticography (ECoG) surgery

Two adult male Wistar rats (280–340 g; 9–12 weeks old) with clean external ears were used in this study. All surgical procedures were performed under sterile conditions to implant the headpost and ECoG array. Following the procedure described in previous studies [41–44], anesthesia was induced with pentobarbital sodium (40 mg/kg, intraperitoneal) and atropine sulfate (0.05 mg/kg, subcutaneous), administered 15 minutes before surgery to reduce tracheal secretions. To minimize local discomfort, 2% xylocaine was applied liberally to the incision site. A head-fixation bar was mounted onto the skull using dental cement and six titanium screws. A craniotomy was then performed over the left auditory cortex, exposing an area of approximately 4.5 × 5 mm while keeping the dura mater intact. A 32-channel ECoG array (KD-PIFE, KedouBC, China) arranged in a 6 × 6 grid with 1 mm interelectrode spacing was implanted, covering roughly a 5 × 5 mm region encompassing the entire auditory cortex (AC) (Fig. 4A).

Following implantation, the array was protected with artificial brain gel, the bone flap was repositioned, and the site was sealed with dental cement to ensure closure and prevent infection. Postoperative care lasted seven days, during which animals received anti-edema medication and antibiotics. Body weight was monitored daily to ensure normal recovery. All animal procedures were approved by the Animal Subjects Ethics Committee of Zhejiang University (approval number: ZJU20210078).

#### Extracellular recording surgery

Two adult male Wistar rats (280–340 g; 9–12 weeks old) were used for extracellular electrode recordings. The surgical preparation followed procedures similar to those used for ECoG implantation to ensure stable head fixation. After surgery, the left lateral and dorsal surfaces of the skull were exposed. Once cleaned and dried, reference landmarks were marked at the bregma and on the left lateral skull to facilitate precise electrode placement.

Recordings targeted the left primary auditory cortex. To prevent tissue overgrowth, the skull overlying the target region was lightly coated with dental cement and enclosed within a protective wall. A small craniotomy (∼2 × 2 mm) was created at the designated site using a handheld micro-drill and subsequently sealed with brain gel. On the recording day, the dura mater was gently punctured, and the recording electrode was vertically advanced into the target region under the guidance of the previously established landmarks.

#### Auditory stimuli and Experimental Procedures

Three experimental paradigms were developed for the EEG, ECoG and extracellular recording experiments, all employing transitional click trains composed of 200-µs pulses. A transitional click train was generated by concatenating two regular click trains. For instance, a Reg_4-4.06_ transitional train consisted of a regular click train with an inter-click interval (ICI) of 4 ms, immediately followed by another with an ICI of 4.06 ms. For continuous click trains that transitioned seamlessly from ICI□ to ICI□, the transition time was defined as the onset of the first click following the first ICI□ interval (Fig. 2A). The difference between the two ICIs was quantified by the ratio of ICI□ to ICI□. The sound intensity of all click trains was calibrated to 60 dB sound pressure level (SPL) to ensure consistent stimulation across both human and animal experiments. Calibration was carried out using a ¼-inch condenser microphone (Brüel & Kjær 4954, Nærum, Denmark) connected to a PHOTON/RT analyzer (Brüel & Kjær, Nærum, Denmark). We also recorded the sound pressure of the stimuli used in the free-field experiments (human Experiment 1 and the rat ECoG experiment; Fig. 1) using a microphone with a sensitivity of 2.86 mV/Pa. The recorded signals were subsequently converted to sound pressure level by calculating time-resolved RMS amplitudes (50-ms bin, 1-ms step), expressed in decibels relative to 20 μPa.

For human EEG recordings, click stimuli were generated in MATLAB R2022b (The MathWorks, Inc., Natick, MA, USA) on a computer equipped with a high-fidelity sound card (Creative AE-7). The clicks were synthesized at a sampling rate of 384 kHz with 32-bit precision and presented via a stereo speaker system (Golden Field M23), ensuring precise temporal delivery of the click train sequences. In contrast, rat ECoG and extracellular recording experiments were performed in a soundproof room, with stimuli delivered contralaterally to the recording site in rats. Acoustic stimuli were digitally generated using a computer-controlled auditory workstation (RZ6, TDT) at a sampling rate of 97656 Hz and presented through magnetic speakers (MF1, TDT) integrated within the TDT system.

#### EEG experimental procedures

The EEG study comprised three change detection experiments. Each stimulus was repeated a minimum of 40 times to each participant across all EEG experiments.

##### Experiment 1 (ICI contrast, Fig. 2A)

In this experiment, participants were positioned in a chair facing a keyboard with earphone wearing. This experiment included 15 regular transitional trains consisting of 3 simulated echo levels and 5 interval contrasts.

The room acoustics were simulated using the *Roomsimove toolbox* for MATLAB [45], which is based on the image source method (ISM). The ISM models echo by creating virtual image sources for each wall reflection, and the resulting room impulse response represents how sound propagates and reverberates in the room. The simulated room measured [3.63, 2.09, 2.6] m, with the sound source located at [1.28, 0, 0.8] m and the receiver at [1.68, 0.4, 1.3] m, all of these parameters are determined according to the real measurement. The six simulated room surfaces were assigned frequency-dependent absorption coefficients: Level 0 (no processing), Level 1 = 0.95, and Level 2 = 0.4, representing different acoustic environments from highly absorptive to more echoic conditions.

In each echo level, an initial 1-s regular click train with an inter-click-interval (ICI) of 4 ms was played, followed by another 1-s regular click train with ICIs randomly selected from 5 potential contrasts: 0, 0.25%, 0.5%, 0.75%, and 1.5%, corresponding to the intervals of 4, 4.01, 4.02, 4.03 and 4.06 ms. Participants were required to report whether the sound changed by pressing two designated keys within 2 seconds after the end of the stimuli, with right key representing change in the transitional stimulation and left key for no change (Fig. 2A).

##### Experiment 2 (Interval insertion, Fig. 2D)

In this experiment, participants listened to the stimuli through earphones under three echo-level conditions. For each level, 2, 8, or 32 intervals—each corresponding to a 4.06-ms ICI and containing no silent gaps—were inserted into a regular click train with a 4-ms ICI (Fig. 2D). These inserted intervals represented slightly lengthened ICIs rather than silent pauses, thereby maintaining continuous acoustic stimulation while modifying the temporal pattern of the sound. The control conditions included Reg_4-4_ and Reg_4-4.06_.

##### Experiment 3 (Free field, Fig. 2F)

In this experiment, participants listened to auditory stimuli presented from a speaker positioned directly in front of them. Two distinct acoustic environments were established to manipulate sound echo characteristics. In the low-reverb condition, the experiment took place in a soundproof room lined with sound-insulation cotton. In the high-reverb condition, participants were surrounded by a circular enclosure of plastic curtains, creating an echoic environment. The auditory stimuli consisted of transitional click trains identical to those used in Experiment 1, maintaining the same ICI contrast configuration without modification.

#### ECoG experimental procedures

The ECoG experiment followed a design similar to Experiment 3 in the human EEG study, but used different ICI contrasts for the five transitional click trains: Reg_4–4_, Reg_4–4.4_, Reg_4–4.6_, Reg_4–4.8_, and Reg_4–5_. The two echoic environments were also configured differently. A soundproof room lined with acoustic foam was used to create a low-reverb environment, while a corrugated cardboard box mounted on the experimental platform was used to construct a high-reverb environment.

#### Extracellular recording experimental procedures

The study included two sessions specifically designed for extracellular recordings.

##### Session 1 Frequency screening procedure

A sequence of pure tones was randomly presented to the contralateral side of the recording site. Each tone had a duration of 100 ms with 5-ms rise and fall times. The frequency range extended from 0.5 kHz to 48 kHz, divided into 26 logarithmically spaced steps, and sound intensity varied from 0 to 70 dB SPL in 10-dB increments. Each frequency–intensity combination was repeated five times, with an interstimulus interval of 300 ms. This session was designed to characterize basic neuronal response properties, including the frequency response area (FRA) and characteristic frequency (CF).

##### Session 2 Transitional click train procedure

This session adopted the stimulation design of Experiment 1 from the human EEG study, but with different ICI contrasts across five transitional click trains: Reg_4–4_, Reg_4–4.4_, Reg_4–4.6_, Reg_4–4.8_, and Reg_4–5_. The three echo levels were simulated using the same room parameters as in the EEG setup.

### Data collection and pre-processing

#### EEG recording

EEG recordings were obtained using a 32-channel electrode cap (10–20 system; NeuSen W series, Neuracle, China) connected to a wireless amplifier of the same series. Two electrodes placed slightly posterior and anterior to Fz served as the reference (REF) and ground (GND) electrodes, respectively. EEG signals and trial onset markers were recorded simultaneously at a sampling rate of 1 kHz. Participants were comfortably seated with head support in an acoustically isolated room throughout the recording sessions and were instructed to minimize eye blinks and facial movements to reduce artifacts.

EEG preprocessing followed standard procedures. Raw EEG signals were band-pass filtered between 0.5 and 40 Hz and epoched relative to stimulus onset, covering –500 to +1000 ms. Four-second epochs (–1 to +3 s relative to trial onset) were then extracted, and independent component analysis (ICA) [46] was applied to remove electrooculogram (EOG) artifacts. After artifact correction, baseline normalization was performed by subtracting the mean activity during the –200 to 0 ms pre-stimulus window for each trial. Motion artifacts were identified using a relative thresholding method. A data point was labeled as a “bad sample” if its amplitude exceeded the range of mean ± 3 × SD across channels within a trial. The proportion of bad samples determined the relative threshold for trial rejection. Trials containing more than 20% bad samples were classified as “bad trials,” and channels containing more than 10% bad trials were labeled as “bad channels.” Bad channels were excluded first, followed by the removal of bad trials identified from the remaining clean channels. Finally, event-related potentials (ERPs) were computed by averaging the cleaned epochs for each condition, channel, and participant. Before inter-subject analysis, ERP amplitudes were normalized by each channel’s standard deviation within subjects to reduce inter-channel variability.

#### ECoG recording

ECoG recordings were obtained by placing the reference wire from the ECoG array into the subdural space, while the ground wire was connected to a titanium screw fixed to the anterior part of the skull. Lead wires from the array were attached to micro-connectors (ZIF-Clip headstage adapters, Tucker-Davis Technologies, TDT, Alachua, FL). The neural signals were amplified using an RZ5 amplifier (TDT), digitized at a sampling rate of 12 kHz, and stored on hard drives for subsequent offline analysis. The raw ECoG signals were segmented to include 1 second before the onset and 1 second after the completion of each transitional click train. The segmented data were then band-pass filtered between 0.5 and 200 Hz and downsampled to 600 Hz. Baseline correction was applied, and trials or channels containing excessive noise were excluded from further analysis. The preprocessing procedure followed the same methodology as that used for EEG data.

#### Extracellular recording

Extracellular recordings were performed using 128-channel electrodes [47], each penetration was precisely positioned according to predefined anatomical markers. A flexible ground wire was soldered to the electrode’s printed circuit board, with the ground and reference contacts shorted. During the experiments, this ground wire was connected to a titanium screw fixed at the anterior part of the skull. The probes were carefully inserted vertically through 0.3–0.5 mm openings in the dura, avoiding visible blood vessels to minimize cortical damage. Insertion was controlled by a single-axis motorized micromanipulator operated from outside the soundproof room. The average penetration depth within the auditory cortex was 2 ± 0.1 mm (mean ± SD, n = 36) below the cortical surface, consistent with the standard rat brain atlas. The presence of auditory responses was verified using white noise stimulation.

Neural signals were acquired using the Intan RHX Data Acquisition Software (v3.4; Intan Technologies, Los Angeles, CA, USA) at a sampling rate of 30 kHz. Spike sorting was performed offline with the Kilosort 3 algorithm, followed by manual curation in Phy 3. For each neuron, a response window (0–100 ms) and a baseline window (–100–0 ms) were defined relative to the onset of each sound. To assess response reliability, the Poisson cumulative distribution function was computed based on the baseline activity to estimate the likelihood that observed spike counts occurred by chance. The response latency was defined as the first time point at which the Poisson probability dropped below 1 × 10^-3^. Neurons were classified as responsive if they met all three of the following criteria: 1) The firing rate within the response window was significantly higher than that in the baseline window (right-tailed paired t-test, α = 0.05); 2) A response latency could be detected; and 3) The mean spike count during the response window exceeded that during the baseline by at least one spike. Neurons satisfying all criteria were identified as auditory-responsive and included in subsequent analyses. For local field potentials (LFPs), the raw signals were downsampled to 600 Hz for current source density (CSD) analysis.

All AC recordings used for subsequent analyses were obtained exclusively from the left hemisphere, identified based on the tonotopic gradient of CF. In total, 36 penetrations containing 124 neurons were recorded from the AC, and all neurons were included in the final analyses.

#### Data Analysis

All steps of the mentioned off-line EEG, ECoG and extrcellular data processing were performed using MATLAB R2023b and the FieldTrip toolbox [46].

#### EEG data analysis

The onset response and change response were quantified at the EEG electrode P4 across three echo levels in Experiment 1, as shown in Fig. 1C. Time windows were predefined for the onset and change responses: [0, 300] ms for onset and [1050, 1350] ms for change, all relative to the stimulus onset.

To quantify the response at Pz for both the onset and change responses (Fig. 2c and Fig. 3c), the response magnitude (RM) was calculated. The RM is defined as the RMS of the normalized ERP within the predefined time window. Normalization was done by dividing each channel’s ERP by its standard deviation over the entire trial.

The RM calculation followed the formula:

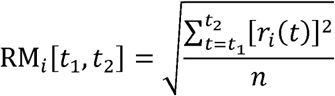

Where *r_i_*(*t*) represents the ERP of the *i^th^* channel at time *t, n* represents the number of samples within the time window from *t*_1_ and *t*_2_ and RM*_i_* is the response magnitude of channel *i*. The RM was computed for each subject and condition within the predefined windows.

#### ECoG data analysis

For each rat, the RM was computed for every channel across all trials. Using the same time window as in the EEG data analysis, amplitudes were measured for both the onset response and the change response. In addition, a differential RM (ΔRM) was calculated, defined as the difference in response magnitude between the high-reverb and low-reverb experimental conditions. Mathematically, it is expressed as:

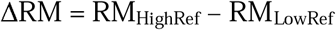

Where RM_HighRef_ is the response magnitude for the high-reverb condition and RM_LowRef_ is the response magnitude for the low-reverb condition.

To evaluate the statistical significance of RM and ΔRM for each ECoG channel, a nonparametric bootstrap procedure with 1000 iterations was performed to generate a null distribution along the trial dimension. Significance was determined by assessing whether zero fell outside the lower bound of the 95% confidence interval (CI) for RM or ΔRM, indicating that the observed values were significantly greater than zero. Channels showing significant RM or ΔRM values were marked with an “x” to illustrate their spatial distribution in Fig. 4F-G and SFigs. 2C-D, 3D-E.

#### Extralcellular data analysis

For spike data, peristimulus time histograms (PSTHs) were generated using a bin width of 30 ms and a step size of 1 ms. For a certain ICI difference *j*, the effect of a certain echo level *i* was defined as

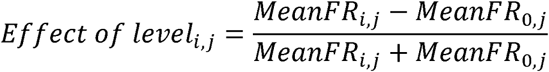

In this formula, MeanFR_i,j_ denotes the mean firing rate from 0 ms to 200 ms after the transition point for ICI difference *j* and echo level *i*.

Since there are two conditions with simulated modification (level 1 and level 2), we constructed a two-dimension space to form a vector describing the global echo effect for each neuron. The vector could provide two characteristics of the echo effect: the vector strength demonstrates the magnitude of echo effect (Fig. 5E), while the phase of the vector indicates the direction of echo effect (Fig. 5F).

To characterize the influence of echo on neuronal sensitivity to ICI contrast, signal detection theory was applied to estimate the neuronal threshold [48–52]. For each echo level and ICI difference, receiver operating characteristic (ROC) analysis was performed using the *perfcurve* function in MATLAB. The analysis compared change responses (200 ms following the change point) with baseline responses (200 ms preceding the change point) and standard responses in firing rate (Fig. 6A, B). The area under the ROC curve (AUC) represented the probability that an ideal observer could discriminate a deviant sound from a standard one based solely on neuronal firing activity. The AUC values were then plotted as a function of the difference between deviant and standard sounds, and the resulting curve was fitted with a normalized Gaussian integral function to quantify neuronal sensitivity [51]:

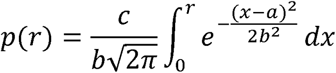

In this expression, ***p*** is the proportion of pressing button choices, ***r*** is the ratio between deviant and standard frequency. The curve fitting procedure was achieved using ‘psignifit’ software package for MATLAB [53]. We also defined the neuronal threshold as the 70 % correct performance of the fitted cumulative Gaussian (Fig. 6C).

#### LFP data analysis

The RM of the LFP was calculated using the same analytical procedure as applied to the EEG data. To visually highlight the differential echo effects between onset and change responses, the RM values were normalized across electrode channels and echo levels relative to the maximum RM (Fig. 7C, D).

To further examine echo effects across cortical layers, we applied the same analytical approach used for the neuronal data. The only modification was that all ICI contrast conditions with nonzero differences were combined to compute the effect vector (Fig. 7E) and vector strength (Fig. 7F).

To identify the onset time at which LFP responses began to diverge across echo levels, a permutation-based analysis of variance (ANOVA) was conducted using the *ft_timelockstatistics* function from the FieldTrip toolbox [46]. At each time point and channel, an F-statistic was computed to compare the three echo conditions. Condition labels were then randomly permuted across trials, and a new *F*-value matrix was generated for each permutation to construct a null distribution at every time point. The observed *F*-values were compared with their corresponding null distributions, and samples with p < 0.05 were considered statistically significant (Fig. 7G). The onset of divergence for each channel was defined as the first time point at which a significant difference emerged (Fig. 7H).

The change detection threshold for the LFP data was calculated using the same method as that applied to the neuronal data.

#### Current Source Density Analysis

One-dimensional CSD profiles were calculated from the second spatial derivative of the LFP signal, as described previously [54–57], and can be approximated by the following formula:

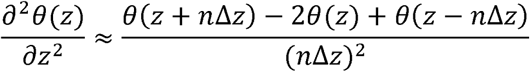

where θ denotes the field potential, *z* represents the spatial coordinate perpendicular to the cortical laminae, Δ*z* is the spatial sampling interval (Δz = 50 µm), and *n* is the differentiation grid (n = 2). To reduce spatial noise, a five-point Hamming filter was applied□. The LFP data used for this analysis were obtained from channels with a 50 µm sampling interval (Fig. 7A). In the resulting laminar CSD profiles, current sinks are shown in red and current sources in blue (Fig. 7B). Visualization of laminar profiles was further enhanced through linear channel interpolation.

In this study, the results from the AC are presented and interpreted with respect to cortical layer organization. The classification of layers was based on two criteria. First, the electrode penetration depth was compared with previously published depth profiles of cortical layers [37, 58]. Second, CSD profiles were analyzed. The latency–depth patterns of CSD sinks in our data closely matched those reported in earlier studies [55–57, 59], f from which we adopted the corresponding layer classification scheme. Based on these two criteria, the cortical structure was divided into three compartments for analysis: supragranular (Sg) layers I–III, granular (Gr) layer IV, and infragranular (Ig) layers V–VI.

#### Statistics

Statistical analyses were performed using several approaches depending on the experimental context.

Two-way repeated-measures ANOVA was used to assess the effects of ICI)contrast and echo level on behavioral performance (Fig. 2C, E) and response magnitudes (RMs) (Fig. 3C-D and Supplementary Fig. 3B-C). Wilcoxon signed-rank tests were applied to compare RMs between high- and low-reverb conditions in the EEG experiments (Figs. 2G and 3E; Supplementary Fig. 1D). Paired t-tests were used to compare the echo effects on onset versus change responses in the ECoG and extracellular recordings (Fig. 4D-E; Fig. 5E; Supplementary Fig. 2A, B; Supplementary Fig. 4B, E, H; Fig. 7F). Friedman tests were employed to evaluate differences across echo levels in experiments involving simulated sounds (Fig. 3C-D; Supplementary Fig. 1B; Supplementary Fig. 3C; Fig. 6D) and to assess layer-dependent variations in the LFP analyses (Fig. 7I; Supplementary Fig. 7C, F, I; Supplementary Fig. 8C). Circular data were analyzed using the *CircStat* MATLAB toolbox [60]. Differences between onset and change responses were evaluated using a paired circular test, implemented as a Rayleigh test on the pairwise angular differences between the two conditions (Fig. 5F and Supplementary Fig. 4C, F, I). For ECoG data, a nonparametric bootstrap test was used to determine the significance of RM and ΔRM for each recording site (Fig. 4F, G; Supplementary Fig. 2C, D; Supplementary Fig. 3D, E). Additionally, a permutation-based ANOVA (1,000 iterations) with FDR correction was performed to identify the earliest time point at which LFP responses diverged significantly across echo levels (Fig. 7G, H; Supplementary Fig. 7A, B, D, E, G, H).

## Supporting information

Supplementary sounds

## Author Contributions

Conceptualization, X.Y.; Methodology, X.Y., P.S., H.X. and H.Y.; Investigation and Data Curation, X.Y., P.S., L.Z., H.X., H.Y., Y.Z., X.B., I.M., N.S.P., Z.T., P.C., T.Z. and X.Z.; Writing – Original Draft, X.Y., P.S., L.Z., H.X. and H.Y.; Writing – Review & Editing, X.Y., X.Z., P.S., L.Z., Y.W., H.X. and H.Y.; Funding Acquisition, X.Y., X.Z. and Y.Z.; Resources, X.Y.; Software, P.S., H.X. and H.Y.; Validation and Visualization, P.S., H.X. and H.Y.; Supervision, X.Y.

## Acknowledgments

We are grateful to Profs. Lin Qing and Xi Chen for their invaluable comments on the early version of the manuscript, as well as to Xiaokai Kou for his assistance with the experiments. This work was supported by STI2030-Major Projects (2022ZD0204600 and 2022ZD0204800) (to X.Y.); National Natural Science Foundation of China 32571216 and 32171044 (to X.Y.) and 32100827 (to Y.Z.); Key Support Discipline Construction Project of Shanghai Municipal Health Commission 2023ZDFC0203 (to X.Z.);

---

**Supplementary Figure 1.**
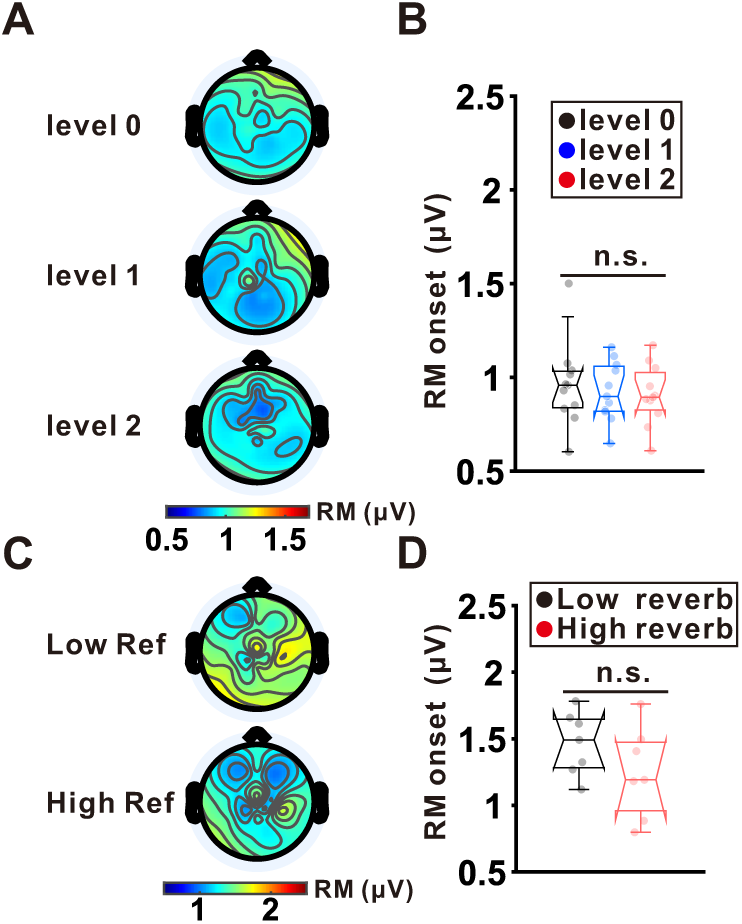
The effect of echo is not obvious for onset response. **(A)** Topographic maps show onset RM distributions under three echo conditions across the scalp in Experiment 1. **(B)** Boxplots show group-level RM values (averaged across electrodes) under level 0 (black), level 1(blue) and level 2 (red) conditions (mean: central line; standard error: box edges; 10^th^-90^th^ percentiles: whiskers). Transparent dots represent individual participants. **(C)** Topographic maps show onset RM distributions under three echo conditions across the scalp in Experiment 3. **(D)** Boxplots show group-level RM values (averaged across electrodes) under low-reverb (blue) environment and high-reverb (red) environment (mean: central line; standard error: box edges; 10^th^-90^th^ percentiles: whiskers). Transparent dots represent individual participants.

**Supplementary Figure 2.**
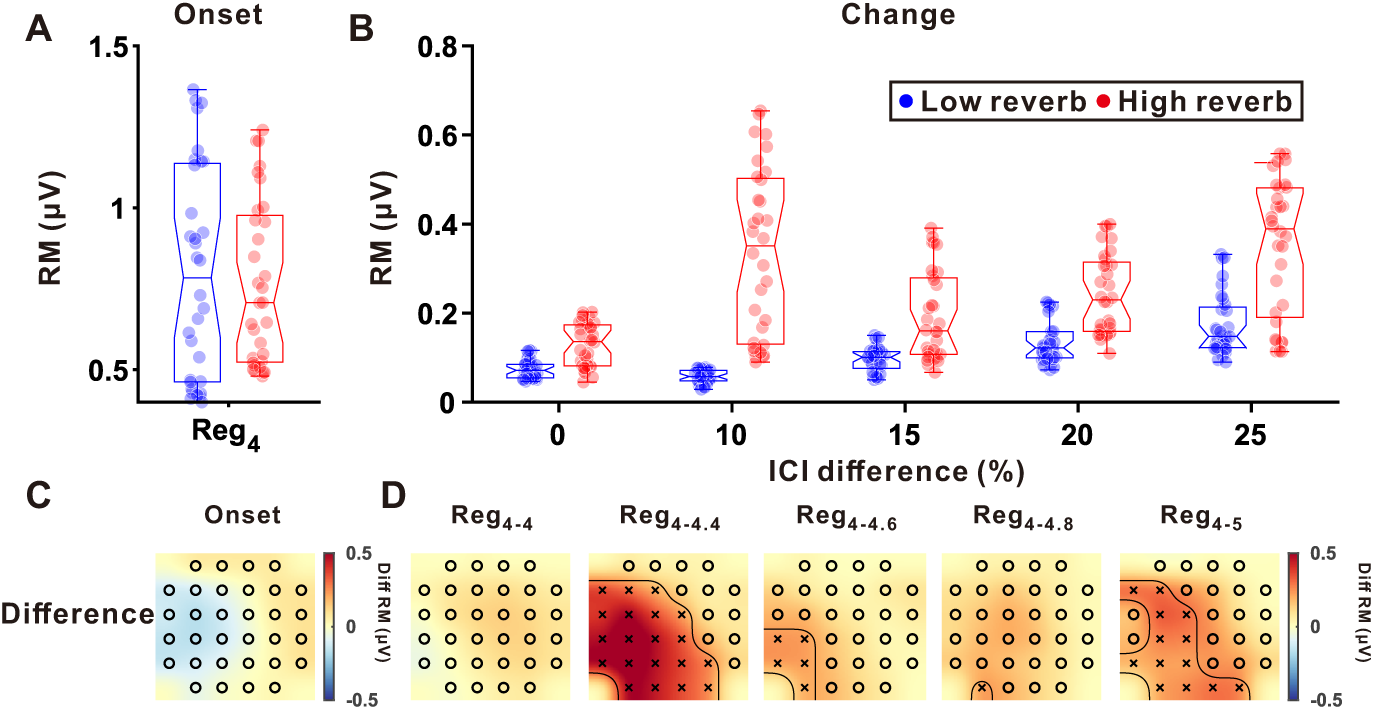
ECoG results from another example rat in two echo environments. **(A)** Boxplots show RM values of onset responses under low-reverb (blue) environment and high-reverb (red) environment (mean: central line; standard error: box edges; 10^th^-90^th^ percentiles: whiskers). **(B)** Boxplots show RM values of change responses under low-reverb (blue) environment and high-reverb (red) environment (mean: central line; standard error: box edges; 10^th^-90^th^ percentiles: whiskers) across five ICI differences. **(C)** Topographic maps show the distribution of onset ΔRM. Channels with significant ΔRM were highlighted with ‘x’ (p < 0.05, nonparametric bootstrap procedure). **(D)** Topographic maps show the distribution of change ΔRM across different ICI contrasts. Channels with significant ΔRM were highlighted with ‘x’ (p < 0.05, nonparametric bootstrap procedure).

**Supplementary Figure 3.**
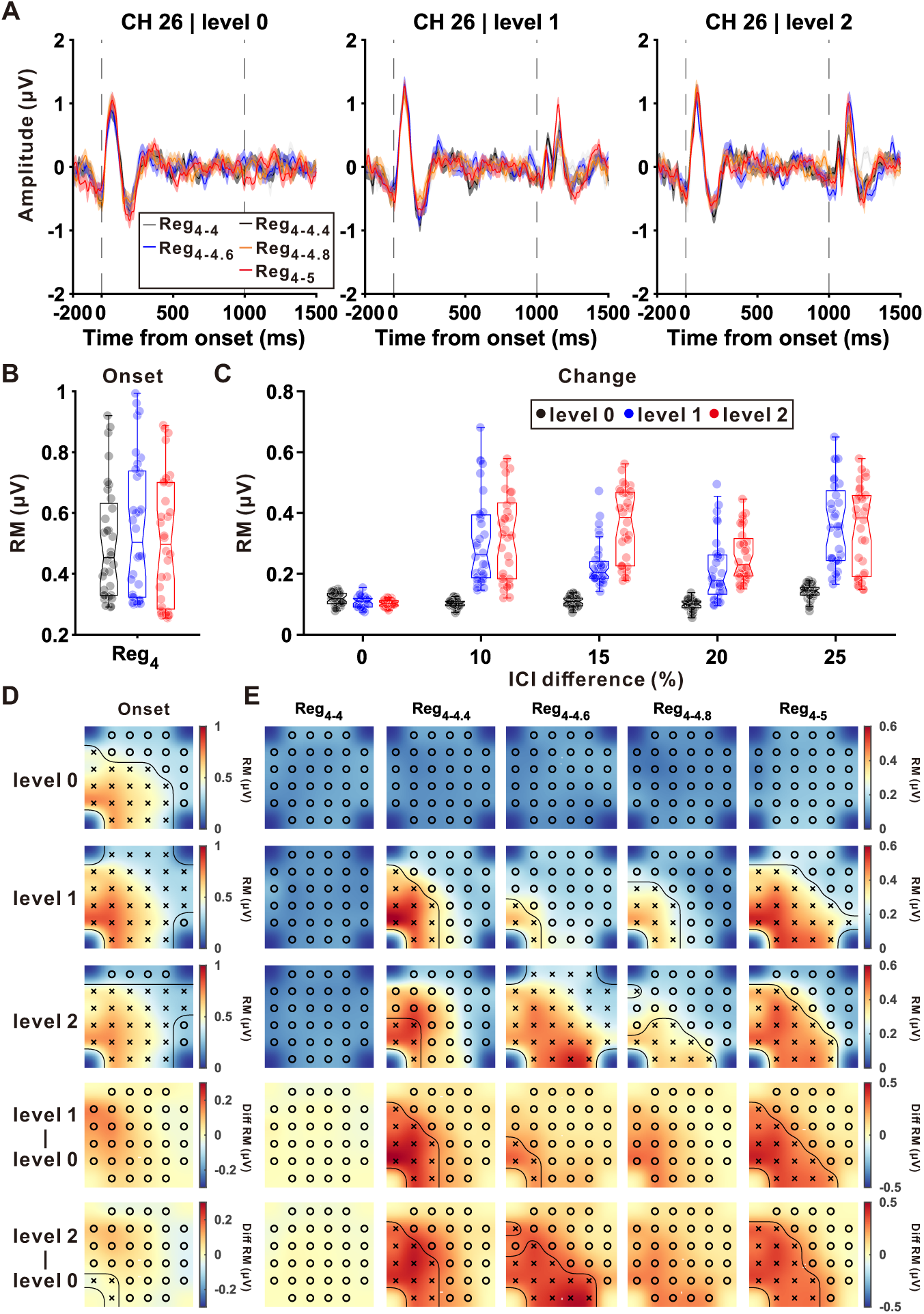
ECoG results from another example rat in response to three simulated echo levels. **(A)** From left to right: Averaged responses at channel 26 across repeated trials under level 0 (left), level 1 (middle) and level 2 (right). Five curves corresponding to different ICI contrasts were colored differently: Reg_4-4_ (gray), Reg_4-4.4_ (black), Reg_4-4.6_ (blue), Reg_4-4.8_ (orange), and Reg_4-5_ (red). The shaded area around the curve represents ± SEM. Vertical black dashed lines represent the onset (0) and change (1000) of the transitional click train, respectively. **(B)** Boxplots (mean: central line; standard error: box edges; 10^th^-90^th^ percentiles: whiskers) show RM values of onset responses across level 0 (black), level 1 (blue) and level 2 (red). **(C)** Boxplots show RM values of change responses across level 0 (black), level 1 (blue) and level 2 (red) across five ICI differences. **(D)** Topographic maps show the distribution of onset RM (first row: level 0, second row: level 1; third row: level 2) and ΔRM (fourth row: difference between level 1 and level 0; fifth row: difference between level 2 and level 0). Channels with significant RM and ΔRM were highlighted with ‘x’ (p < 0.05, nonparametric bootstrap procedure). **(G)** Topographic maps show the distribution of change RM across different ICI contrasts. Channels with significant RM and ΔRM were highlighted with ‘x’ (p < 0.05, nonparametric bootstrap procedure).

**Supplementary Figure 4.**
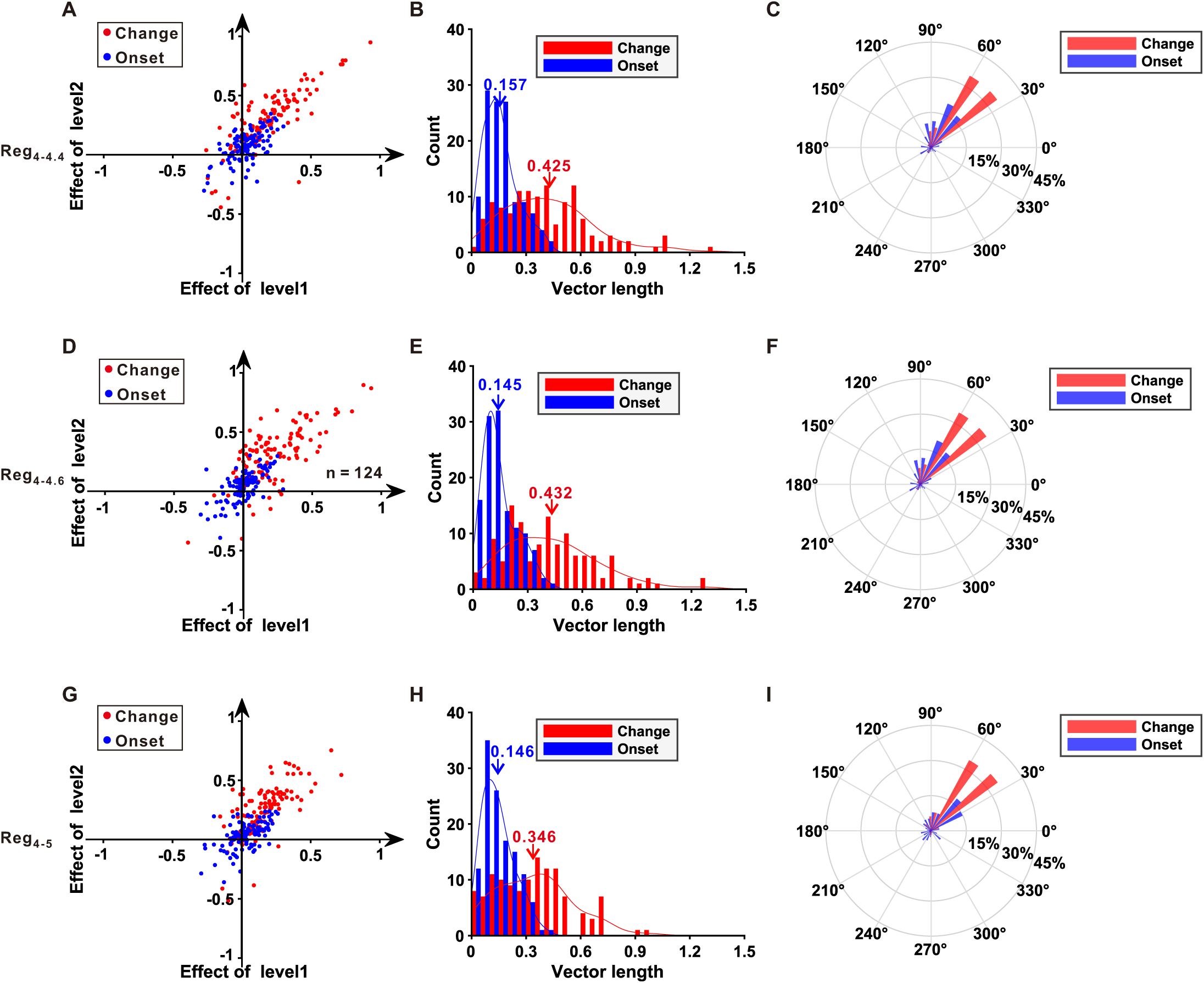
The echo effect for transitional click trains with other ICI contrasts. **(A-C)** Same result representation as D-F in Figure 3, but for Reg_4-4.4_. **(D-F)** The echo effect for Reg_4-4.6_. **(D-F)** The echo effect for Reg_4-5_.

**Supplementary Figure 5.**
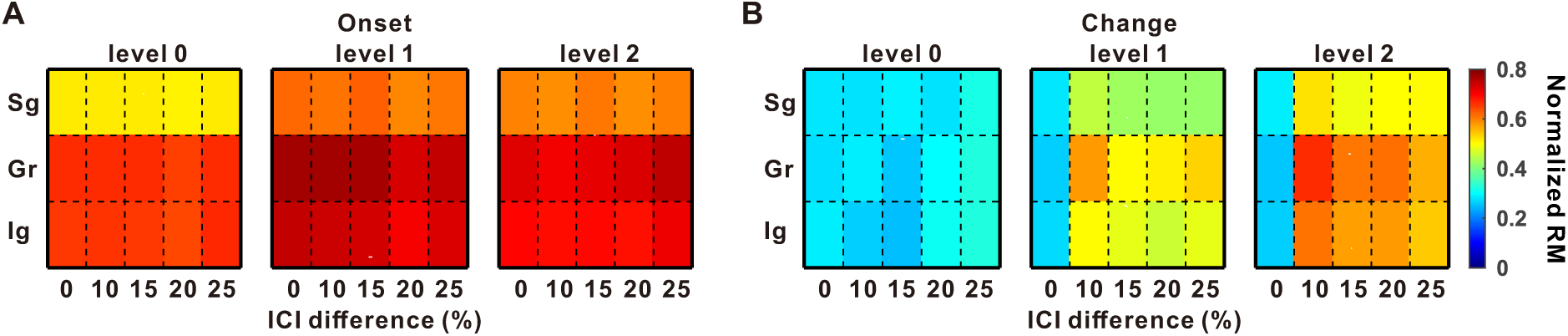
The echo effect on onset and change responses averaged across all penetrations. **(A)** Heat maps showing the averaged normalized RMs across 36 penetrations from two rats for onset responses across three laminar regions (Sg, Gr, and Ig). Within each matrix, RMs are plotted as a function of ICI difference (%), and across matrices as a function of echo level (levels 0–2). **(B)** Same as in (A), but for change responses.

**Supplementary Figure 6.**
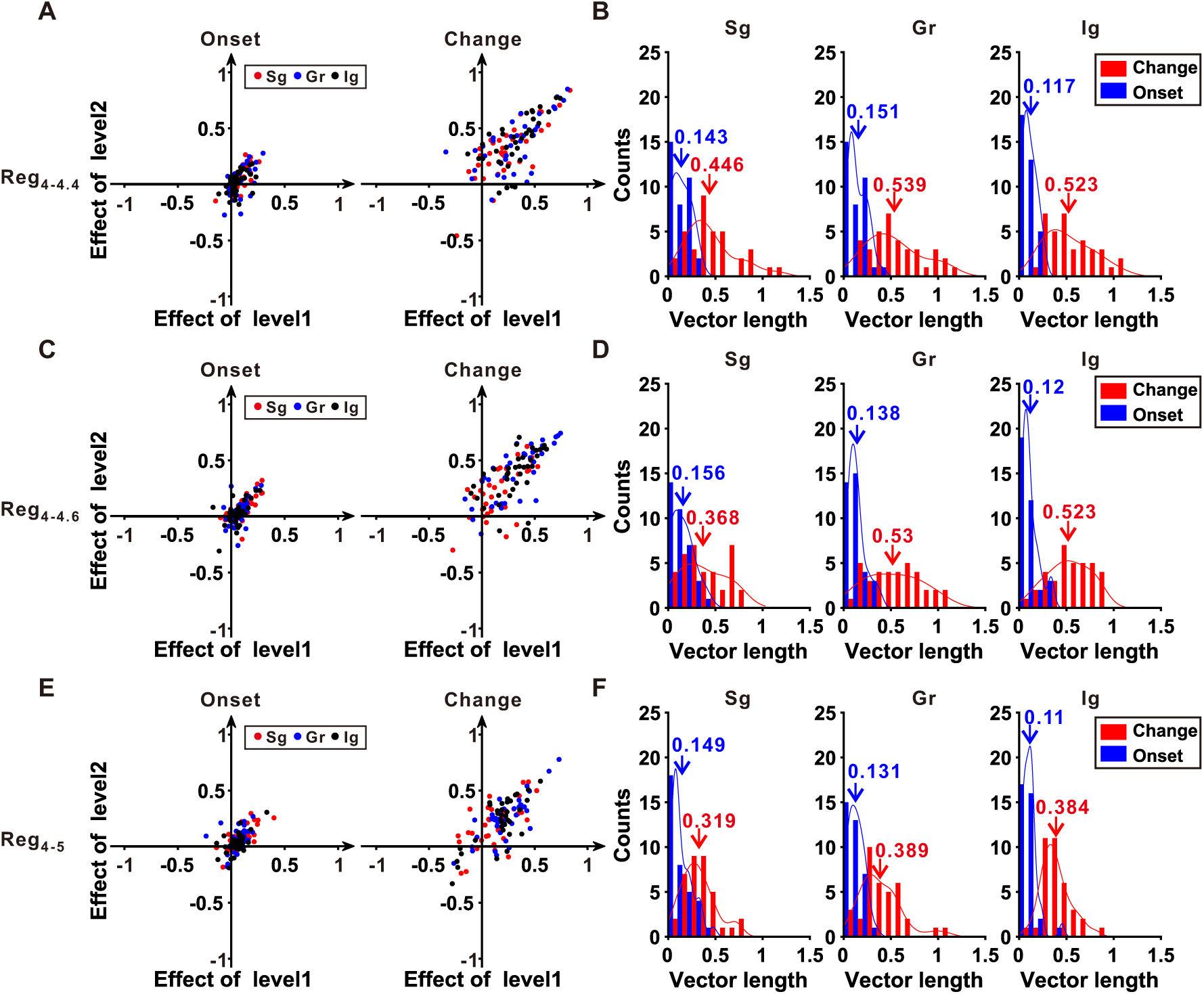
The laminar echo effect for transitional click trains with other ICI contrasts. **(A-B)** Same result representation as E-F in Figure 5, but for Reg_4-4.4_. **(C-D)** The echo effect for Reg_4-4.6_. **(E-F)** The echo effect for Reg_4-5_.

**Supplementary Figure 7.**
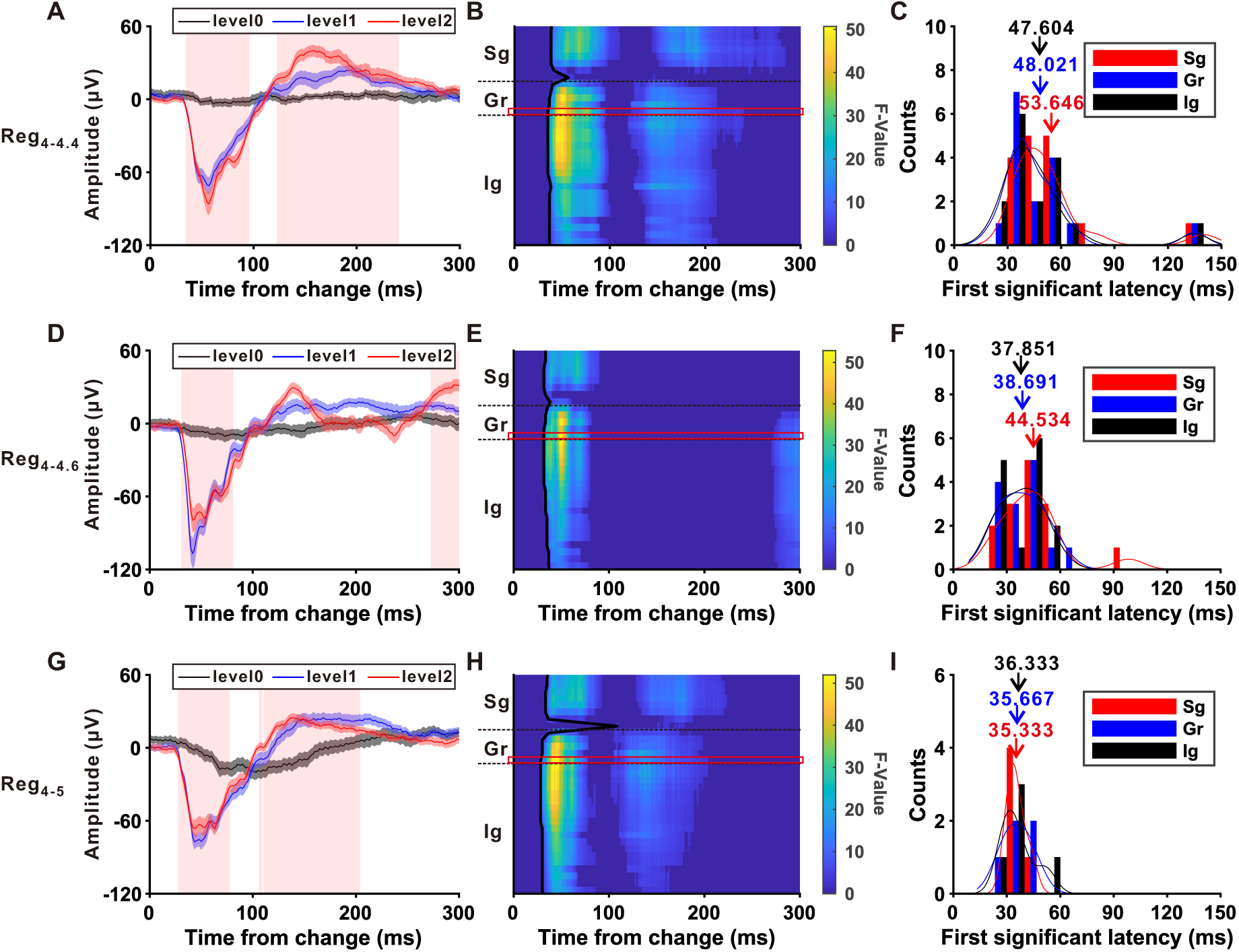
The first significant time for other three ICI contrasts. **(A-C)** Same result representation as A-C in Figure 4, but for Reg_4-4.4_. **(D-F)** The first significant time for Reg_4-4.6_. **(G-I)** The first significant time for Reg_4-5_.

**Supplementary Figure 8:**
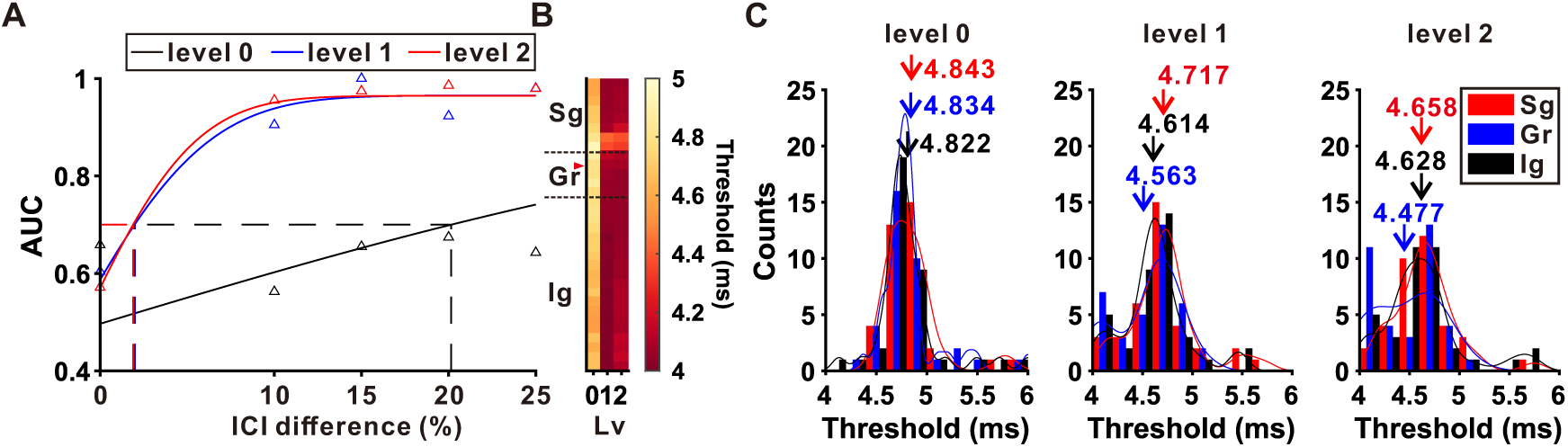

